# Predicting Pain Using a Data-Driven Agent-Based Model of the Bilateral Central Amygdala

**DOI:** 10.64898/2026.04.24.720694

**Authors:** Blesson K Paul, Megan Kwiatkowski, Fatima Zhantibiyeva, Sudhuman Singh, Claire Nguyen, Sri Vaishnavi Devi Singari, Chris Davis Perumal, Iniya Anandan, Anisha P. Adke, Katelyn E. Sadler, Yarimar Carrasquillo, Rachael Miller Neilan, Benedict J. Kolber

**Affiliations:** Department of Neuroscience and Center for Advanced Pain Studies, University of Texas at Dallas, Richardson, Texas 75080; Department of Mathematics and Computer Science, Duquesne University, Pittsburgh, Pennsylvania 15282; National Center for Complementary and Integrative Health, National Institutes of Health, Bethesda, MD 20892; National Institute on Drug Abuse, National Institutes of Health, Bethesda, MD 20892

**Keywords:** Amygdala, Pain, Central Amygdala, Electrophysiology, Neuronal Physiology, Synaptic Plasticity, Calcitonin Gene Related Peptide Receptor, Protein Kinase C δ, Computational Neuroscience, Agent-Based Modeling

## Abstract

Sensory processing in the amygdala is a complex, dynamic process. Decades of surgical, electrical, pharmacological, optogenetic, and chemogenetic *in vivo* manipulations have revealed the nociceptive functions of anatomically- and genetically-restricted neuronal populations. In parallel, molecular and electrophysiological approaches have allowed for high-resolution, temporal examination of nociceptive-induced alterations in amygdala plasticity. Computational integration of this data is critical for future therapeutic development; in practice, these models would allow for *in silico* prediction of amygdala activity following injury, and in a reciprocal fashion, changes in pain-like behaviors following manipulation of discrete amygdala neuronal populations. To this end, we developed a three-dimensional computer model of the bilateral central nucleus of the amygdala (CeA). We employed agent-based modelling to integrate wet-lab data from two CeA cell populations: Calcitonin Gene-Related Peptide Receptor (CGRPR; *Calcrl*) expressing cells and Protein Kinase C delta (PKCδ; *Prkcd*) expressing cells. We integrated the spatial location, connectivity, neuronal activity, and electrophysiological properties of these neurons in our realistic bilateral model architecture. Our model captures properties of the amygdala that drive pain modulation, including hemisphere-specific physiological differences, and generates predictions of nociception related to bladder injury. Predictions from the model were compared retrospectively to pain outcomes during manipulation of CGRPR-expressing neurons in whole mice. These comparisons show strong alignment between our model and *in vivo* outcomes.

**Significance Statement:** Nociceptive processing in the amygdala emerges from interactions among cell types. No framework has been developed that recapitulates this complexity. Here, we generated a 3-D agent-based computational model of the bilateral central amygdala (CeA) using data that accounts for pain-associated changes in neuronal physiology, cellular identity, anatomical positioning, and temporal specificity. The model revealed that hemispheric differences in intrinsic CeA neuronal excitability are dominant drivers of nociceptive output. Furthermore, when manipulated to mimic injury-associated plasticity, the model accurately predicted behavioral outcomes. This model is a publicly available computational tool that predicts hemisphere-targeted pain therapy efficacy. Moreover, the flexibility of this framework will allow future adaptations to other brain areas and disease contexts.

## Introduction

The central nucleus of the amygdala (CeA) is a critical hub in supraspinal nociceptive networks. With dual roles as the primary output nucleus of the amygdala and a recipient of both crude and processed nociceptive inputs from its parabrachial (PBN) (1-3) and basolateral amygdala (BLA) (4, 5) connections respectively, the CeA is a critical inflection point in nociceptive arcs. Activity of CeA neurons is affected by peripheral inputs in a “bottom-up” fashion and, in a reciprocal “top-down” manner, manipulation of these same neurons modulates pain-like behaviors (6, 7). Maladaptive CeA plasticity is widely reported in both patients (8) and animal models with chronic pain (1, 9-11). In particular, significant attention has been given to the CeA in the right hemisphere; neurons in the right CeA frequently exhibit hyperactivation or hyperexcitability in persistent pain states (7, 11-14), and non-specific inhibition of this activity routinely decreases pain-like behaviors in animal models (11, 15).

Although important foundational observations, these early lateralized experiments did not account for additional biological attributes that may ultimately have a tremendous impact on CeA-centered therapeutic efficacy. For example, scRNA sequencing studies have now made the molecular heterogeneity of CeA neurons abundantly clear (16, 17). Most recent estimates propose that more than 25 distinct populations exist in a region that was once described as containing “only GABAergic neurons” (18). Despite not knowing the exact function of each population, initial studies have started to reveal divergent nociceptive properties. For example, in the right CeA, activation of protein kinase Cδ (PKCδ)-expressing neurons can induce pain in uninjured animals whereas activation of somatostatin (SST) expressing neurons reduces pain in animals with neuropathic injury (19). To make matters more confusing, the same neuronal population can also have opposing nociceptive functions depending on hemispheric localization. For example, activation of calcitonin gene related peptide (CGRP) receptor (CGRPR)-expressing neurons in the left CeA decreases pain-like behaviors while activation of the same neuronal population in the right CeA increases pain-like behaviors (11, 15). Each molecular class can be further subdivided by physiological parameters. For example, CGRPR neurons can be sub-classified as regular spiking, late firing, or spontaneously active based on whole cell patch clamp recordings (11); the nociceptive importance of these physiological distinctions remains unclear. And lastly, the temporal contributions of each cell populations are unclear at acute and chronic phases of pain. However, only recently studies have been exploring the differential physiology at acute and chronic pain states on populations like corticotrophin-releasing factor (CRF) expressing and CGRPR-expressing neurons (1, 11).

Given the increasing number of variables that factor into CeA nociceptive status and the accumulating data associated with each state, computational integration of different variables is now needed to accurately predict therapeutic efficacy and prioritize future wet lab experiments. To this end, we developed a three-dimensional (3-D) agent-based model (ABM) of the left and right CeA (20-25). Agent-based modeling is a computational approach for simulating complex systems in which individual “agents” (here, neurons) act and interact with neighboring agents according to programmed set of rules, such as firing rate, spatial location, and dendritic arborization size. We collected anatomical, molecular, and temporally registered physiological measures from CeA neurons under naïve conditions and following cyclophosphamide (CYP)-induced bladder pain. Beginning with CGRPR and PKCδ-expressing populations, data were used to develop a 3-D ABM of the bilateral CeA and emergent nociceptive output. We report that our bilateral ABM revealed that asymmetry in intrinsic excitability between the left and right CeA, rather than upstream molecular differences, is a predominant driver of emergent CeA nociceptive output. Also, our ABM accurately predicted the pain outcomes of *in vivo* manipulations of CGRPR-expressing neurons.

## Results

### Neuronal excitability of CGRPR-expressing cells increased in the right CeA in an acute bladder pain model

To develop a meaningful model of the left and right CeA in pain states, we began by measuring the physiological properties of CGRPR and PKCδ-expressing neurons. These two populations were selected because they are known to receive nociceptive information and undergo excitability changes after injury (11, 19). For the first set of electrophysiological recordings, transgenic mice expressing *Calcrl* promoter-dependent Cre-recombinase were bred with Ai14-tdTomato mice to allow for fluorescent visualization of CGRPR positive neurons (**Figure 1A, 1B)**. Whole-cell patch-clamp recordings were completed on brain slices obtained from mice that underwent cyclophosphamide (CYP)-induced bladder sensitization or that received saline control injections (**Figure 1C**). Recordings were completed 1 day following the final CYP or saline injection, a timepoint that – in CYP-treated mice – reflects an acute inflammatory pain state.

**Figure 1:**
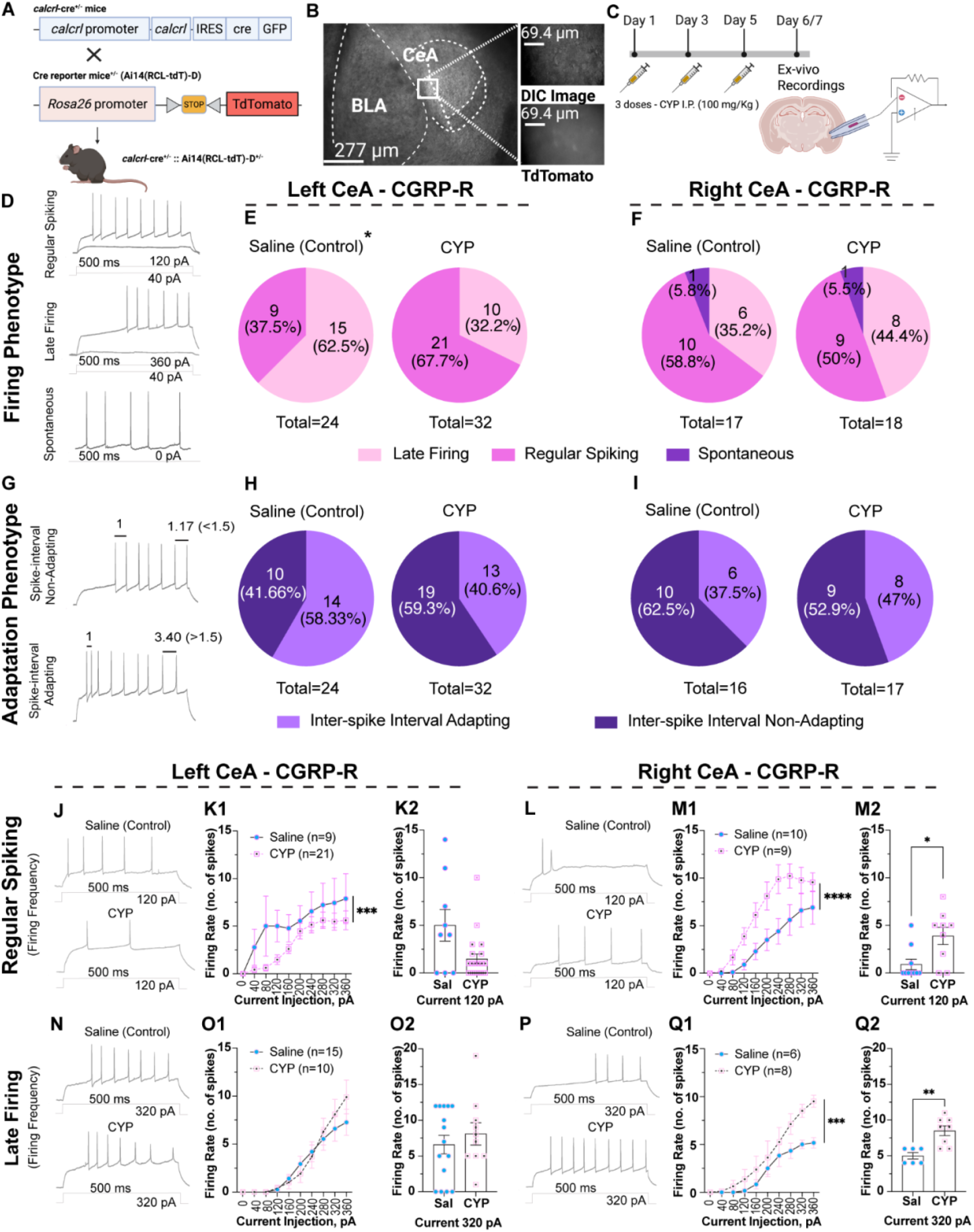
Evaluation of electrophysiological properties of CGRPR-expressing neurons in the left and right CeA. (**A**) Schematic representation of the genetic strategy to target CGRPR-expressing neurons using *calcrl*^cre^ and Ai14 (TdTomato) mice. (**B**) Representative image of tdTomato expression in the CeA with high-magnification insets showing labeled soma. (**D**) Experimental timeline for drug administration (1– 5 DPI) followed by acute slice preparation and whole-cell patch-clamp recordings on 6/7 DPI (**C**). Representative voltage traces of distinct firing patterns observed in recorded CGRPR-expressing neurons, including late-firing (LF), regular-firing (RS), and spontaneously firing phenotypes. (**E, F**) Pie charts illustrate the distribution and percentage of these firing phenotypes under saline and CYP conditions, showing a statistically significant shift (*) in distribution in the left CeA. (**G**) Traces showing firing modalities (non-adapting vs adapting) by their inter-spike interval phenotype. (**H, I**) Pie charts illustrate the distribution and percentage of these adapting phenotypes under saline and CYP conditions with no significant changes observed. **(J, L, N, P)** Representative traces of 500 ms current-evoked spikes in saline (top) and drug-treated (bottom) groups for RS and LF neurons. Summarized input-output curves showing mean firing frequency (Hz) as a function of injected current (pA) for **(K1, M1)** RS type and **(O1, Q1)** LF type. **(K2, M2, O2, Q2)** Adjacent bar graphs quantify the mean firing frequency at specific current steps; individual data points represent single neurons. Data are presented as mean ± SEM. \*\*\*\**P* < 0.0001, \*\*\**P* < 0.001, \*\**P* < 0.01, \**P* < 0.05.

Consistent with prior whole-cell recordings from the CeA-that defined their firing phenotypes (19, 26), we found that CGRPR expressing neurons similarly fell into three categories: regular spiking, late firing, or spontaneously active (**Figure 1D**-). Comparing the saline vs. the acute bladder pain group in the left CeA, we observed a significant difference in firing phenotype prevalence; late-firing neurons were more abundant in the saline group compared to injured group (**Figure 1E)**. Unlike the left CeA, the right CeA did not show any change in the proportion of firing phenotypes under acute bladder pain conditions, except there were spontaneously active neurons observed in addition to regular spiking and late firing neurons (**Figure 1F**).

In addition to firing phenotype, we also categorized the adaptation profile and firing rates of all CGRPR-expressing neurons. Neurons were first classified as either non-adapting (inter-spike interval remained relatively unchanged during sustained depolarization) or adapting (inter-spike interval increased over sustained depolarization; **Figure 1G**). No significant differences in adaptation type were observed between hemispheres or between control/bladder pain conditions (**Figure 1H, I**). Following this, excitability was assessed in both regular spiking (**Figure 1J-1M**) and late firing CGRPR neurons (**Figure 1N-Q**). During acute bladder pain, regular spiking CGRPR neurons exhibited hemisphere-specific changes in excitability; CGRPR neurons in the left CeA exhibited a decrease in firing rate whereas CGRPR neurons in the right CeA exhibited an increase in firing rate. Late firing CGRPR neurons in the right CeA also exhibited increases in excitability during bladder pain; no injury-associated change in excitability was noted in left CeA CGRPR late firing neurons.

Additional measurements of neuronal excitability such as the resting membrane potential, capacitance, rheobase, action potential amplitude, rise time, duration, current threshold, and voltage threshold did not exhibit any change between control and bladder pain conditions in the regular spiking phenotype belonging to the left CeA (**Figure S1 Ai-iii, Bi-iii, Ci-ii**). However, in the right CeA, the rheobase measurement of regular spiking phenotype exhibited a reduction in current needed for evoking one action potential (**Figure S1 Diii**). This indicates an increase in excitability recapitulating the firing rate data in the right CeA regular spiking phenotype. The rest of the excitability measurements in the right CeA with regular spiking phenotype did not show any change between control and bladder pain conditions (**Figure S1 Di-ii, Ei-iii, Fi-ii**). Additional excitability measurements in the late firing phenotype of CGRPR-expressing neurons in the left and right CeA did not exhibit any change in the bladder pain group compared to the control group (**Figure S1 Gi - Lii**).

### Neuronal excitability of PKCδ-expressing cells decreased in the left CeA in an acute bladder pain model

To complement CGRPR-expressing neuronal recordings, identical whole cell patch clamp assessments were also performed on PKCδ-expressing CeA neurons. Transgenic mice expressing *Pkcd*-dependent Cre-recombinase were bred with Ai14-tdTomato mice to visualize PKCδ-positive neurons (**Figure 2A - B)**. Similar to previous publications from the CeA (19, 26), we observed PKCδ-expressing neurons exhibiting regular spiking, late-firing, or spontaneously active firing phenotypes (**Figure 2D**). Unlike CGRPR-expressing neurons, PKCδ-expressing neuronal firing phenotypes were found in similar abundance regardless of hemisphere and bladder pain state (**Figure 2E - 2F**). Adaptation profiles of PKCδ-expressing neurons were also similar between hemispheres and injury states (**Figure 2G - 2I)**. Excitability of regular spiking and late firing PKCδ-expressing neurons was measured. Although no injury-associated differences were reported in left CeA regular spiking PKCδ-expressing neuronal activity (**Figure 2J - 2K**), regular spiking PKCδ-expressing neuron in the right CeA exhibited increased excitability during acute bladder pain (**Figure 2 M1, M2**). Unlike CGRPR-expressing neurons, PKCδ-expressing late firing neurons in both the left and right CeA exhibited decreased excitability during acute bladder pain (**Figure 2 O1, O2, Q1, Q2**). These data, coupled with additional excitability measures (**Figure S2**) once again revealed unique, injury-dependent changes in excitability that were restricted to specific neuronal populations in the CeA. These unique electrophysiological properties from CeA CGRPR and PKCδ-expressing neurons were used for ABM parametrization.

**Figure 2:**
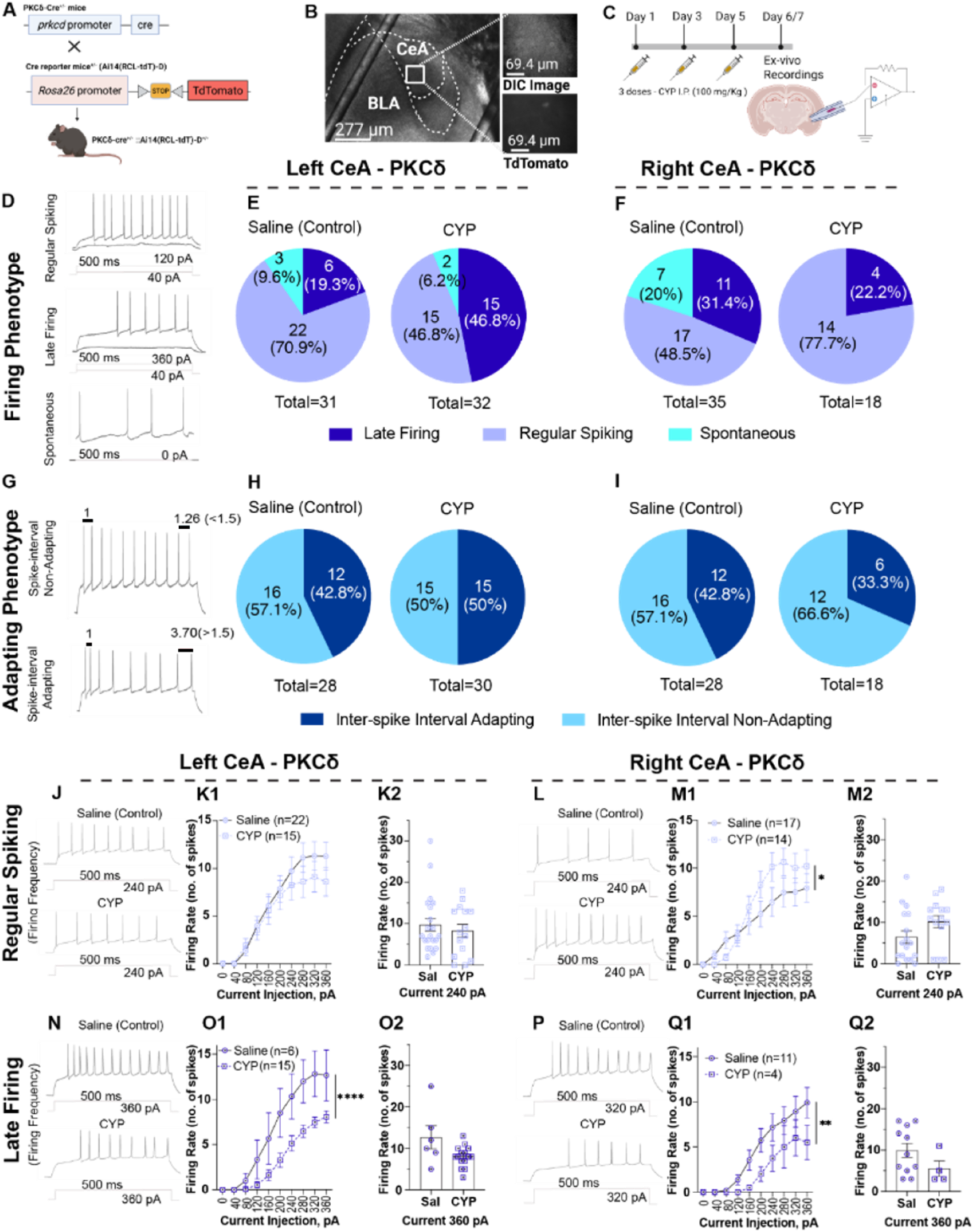
Evaluation of electrophysiological properties of PKCδ-expressing neurons in the left and right CeA. (**A**) Schematic representation of the genetic strategy to target PKCδ-expressing neurons using *prkcd*^cre^ (PKCδ) and Ai14 (TdTomato) mice. (**B**) Representative image of tdTomato expression in the CeA with high-magnification insets showing labeled soma. (**C**) Experimental timeline for drug administration (1– 5 DPI) followed by acute slice preparation and whole-cell patch-clamp recordings on 6/7 DPI. (**D**) Representative voltage traces of distinct firing patterns observed in recorded PKCδ-expressing neurons, including regular-firing, late-firing, and spontaneous/other phenotypes. (**E, F**) Pie charts illustrate the distribution and percentage of these firing phenotypes under saline and CYP conditions in the left and right CeA showing no significant changes. (**G**) Traces showing firing modalities by their inter-spike interval adapting phenotype. (**H, I**) Pie charts illustrate the distribution and percentage of these adapting phenotypes under saline and CYP conditions with no significant changes. (**J, L, N, P**) Representative traces of 500 ms current-evoked spikes in saline (top) and CYP-treated (bottom) groups for RS and LF neurons. Summarized input-output curves showing mean firing frequency (Hz) as a function of injected current (pA) for (**K1, M1**) RS type and (**O1, Q1**) LF type. (**K2, M2, O2, Q2**) Adjacent bar graphs quantify the mean firing frequency at specific current steps; individual data points represent single neurons. Data are presented as mean ± SEM. \*\*\*\**P* < 0.0001, \*\**P* < 0.01, \**P* < 0.05.

### Bladder distention and injury induce ERK phosphorylation in the amygdala

While the firing properties determined by the previous experiments on the CeA population conveys the extent to which they are excitable, molecular markers such as the phosphorylation of extracellular signal related kinase 1/2 (pERK 1/2) conveys the neuronal activation status. pERK 1/2 occurs in the CeA following somatic nociceptive events and is required for the behavioral sensitization that accompanies injury (13, 27). To determine if activation of this protein is elevated in a lateralized manner following visceral bladder pain, Western blot analyses were performed on left and right CeA homogenates obtained at various times following urinary bladder distention in CYP bladder-sensitized animals. Animals’ bladders were distended in triplicate at the following pressures and in the following order: 15, 30, 45, 60, 75 mmHg.

pERK1/2 expression was evaluated via Western blot 5 min following the completion of graded (UBD) in saline and CYP-treated animals. No significant change in pERK1 expression was noted with UBD in either saline or CYP-treated animals (**Figure S3A**; Two-way RM ANOVA, no main effect of distension in saline P=0.2259 or CYP P=0.3161; n=10-11). Additionally, CYP treatment alone did not induce pERK1 expression (Two-way RM ANOVA across all sham animals, no main effect of CYP P=0.6152). Conversely, in both saline and CYP-treated animals, UBD significantly increased ERK2 phosphorylation (**Figure S3B**; Two-way RM ANOVA, main effect of distension in saline p=0.0349, main effect of distension in CYP p=0.0298; n=10-11). Statistically significant increases in this protein were specifically noted in the right CeA of CYP-treated animals that underwent UBD when compared to CYP-treated sham animals (Bonferroni post-test CYP right distend vs. CYP right sham *P<0.05). CYP treatment alone did not increase pERK2 expression in sham animals (Two-way RM ANOVA no main effect of CYP in sham P=0.4596).

### CGRPR and PKCδ Expression data with pERK

To inform the spatial parameterization of the model, we assessed the distribution of CGRPR-expressing, PKCδ-expressing, and pERK-expressing neurons across the anterior-to-posterior extent of the CeA. Specifically, we evaluated marker expression in tissue sections from mice following neuropathic pain injury (“cuff model”). These data were previously published as average values across mice (28). Here, we show the variability that exists in expression values across the eight individual mice included in the original data (**Figure 3A - 3F**). This inter-animal variability includes differences in the number and distribution of CGRPR-expressing and PKCδ-expressing cells along the anterior-to-posterior CeA and variability in pERK activation with cuff injury. Animal-specific parameters were included in the model to examine how biological variability in neuronal expression patterns influences predicted interactions between PKCδ and CGRPR-expressing neurons and emergent nociceptive output.

**Figure 3:**
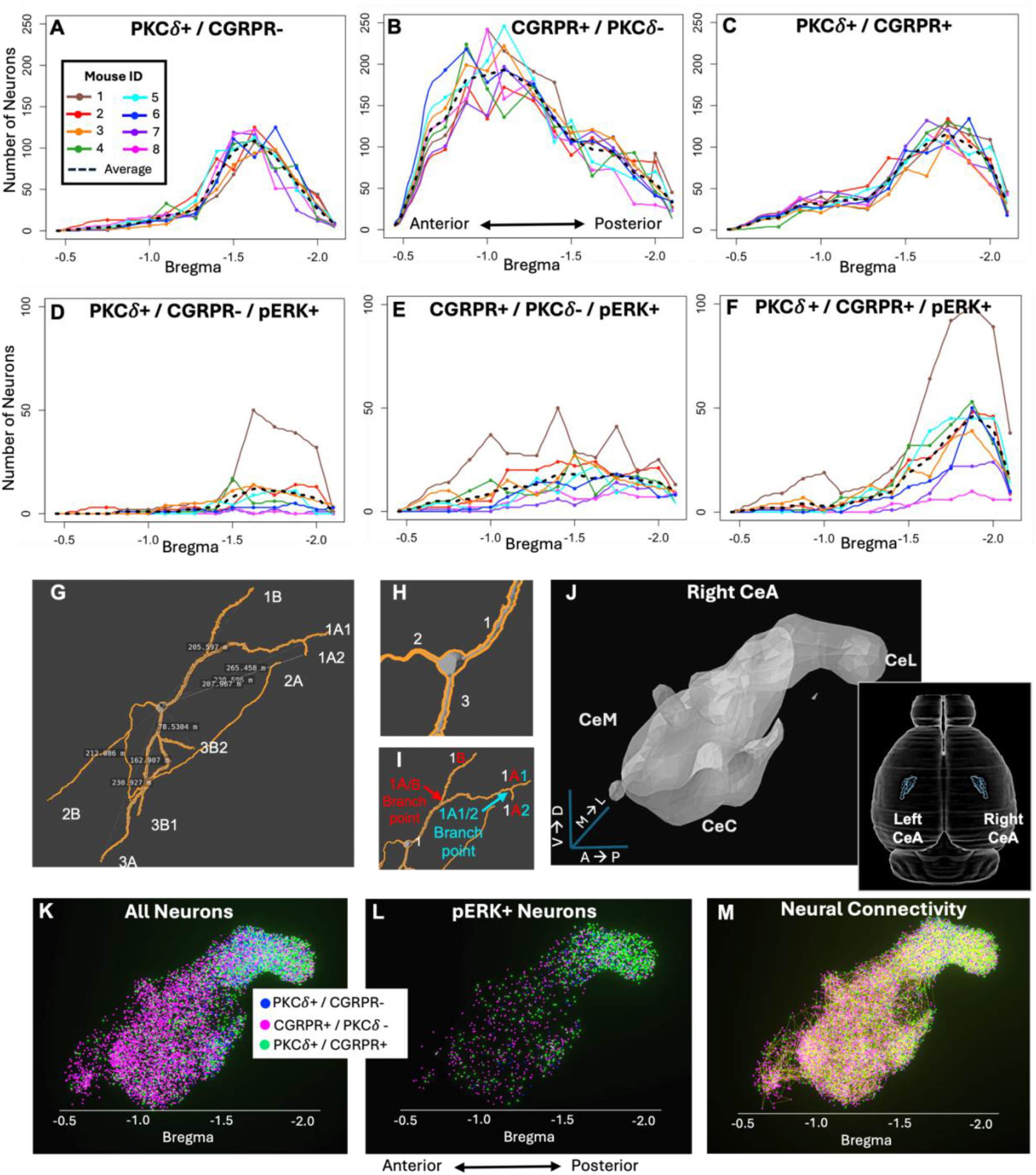
Quantification of PKCδ-, CGRPR-, and pERK-expressing neurons along the anterior-to-posterior (A⟶P) CeA axis. **(A–C)** Distributions of PKCδ, CGRPR, and pERK-expressing neurons along the A⟶P axis from individual mice (n = 8). **(D - F)** Corresponding distributions of pERK expressing neurons. **(G)** A PKC-δ neuron is shown fully labeled and measured. **(H)** Starting at the middle, the dendrite branches are labeled from 1-3, with branch 1 being the longest and branch 3 being the shortest. **(I)** Then the branch point within branch 1 is highlighted, as the first branching point is labeled 1A (longer branch) and 1B (shorter branch). Then, the branch point within branch 1A is further highlighted, as the second branching point is labeled 1A1 (longer branch) and 1A2 (shorter branch). **(J)** 3-D anatomical CeA domain derived from the Blue Brain Cell Atlas. **(K)** Reconstructed 3-D distributions of PKCδ (blue), CGRPR (magenta), and overlapping PKCδ / CGRPR (green) neuron populations for the average mouse (dashed lines in A-C). **(L)** Subset of neurons in **H** that express pERK (dashed lines in D-F). **(M)** Network of neural connections (yellow lines) generated by ABM for neuronal distribution in **H**.

### Evaluation of PKCδ-expressing neuronal dendrite length

Another important consideration when modeling CeA circuitry in 3-D space is the distance over which neurons form connections. Dendritic length constrains the distance over which local inhibitory interactions can occur and therefore provides an empirical basis for modeling distance-dependent connectivity. To estimate potential connectivity distances between neurons, we evaluated biocytin-filled cells from PKCδ reporter mice. These data were previously published but here we performed a higher-resolution 3-D evaluation of the lengths of primary, secondary, and tertiary dendrites from PKCδ-expressing cell bodies (**Figure 3G - 3J**). Over a sample of PKCδ-expressing neurons (n=7), dendritic processes extended from 10 to 270 um from the cell body (**Table S1**) with a log-normal distribution (**Figure S4)**, with approximately half of these being medium range (30-120 um) processes. These findings were incorporated into the computational model to implement distance-dependent connectivity between PKCδ-expressing and CGRPR-expressing neurons.

### ABM results reveal hemispheric asymmetries as key drivers in CeA nociceptive output

Using experimental data from this study and prior work, we constructed a 3-D agent-based model (ABM) of CGRPR and PKCδ-expressing neurons in the left and right CeA. The bilateral ABM is publicly available through our web application (23) and allows for intuitive user-driven experimentation. The model’s spatial domain reproduces the 3-D topology of the left and right CeA as defined by the Blue Brain Cell Atlas (29) (**Figure 3J**). Users can select mouse-specific or average neuronal distributions and expression patterns (**Figure 3K, 3L**), after which a network of neuronal connections is generated (**Figure 3M**). During model simulation, hemisphere-specific neuronal properties, including firing rate and pERK expression, are updated each time step in response to noxious stimulation and an emergent measure of nociceptive output from the bilateral CeA is computed.

In the model, noxious stimulation is implemented as current injection to individual neurons and the resulting emergent bilateral neural activity is quantified as nociceptive output, a system-level proxy for pain-like behavior. Model simulations of constant noxious stimulation (≥120 pA) produced increased nociceptive output during modeled “injury” and sustained elevated output after injury (**Figure 4A**). Variability within a single simulation (black line; **Figure 4A**) reflects stochastic processes inherent to the ABM (e.g., firing rate variability), whereas grey lines represent variability in output across simulations calibrated with different pERK expression levels (±1 mean absolute deviation (MAD) from n=8 mice; **Figure 4D**).

**Figure 4:**
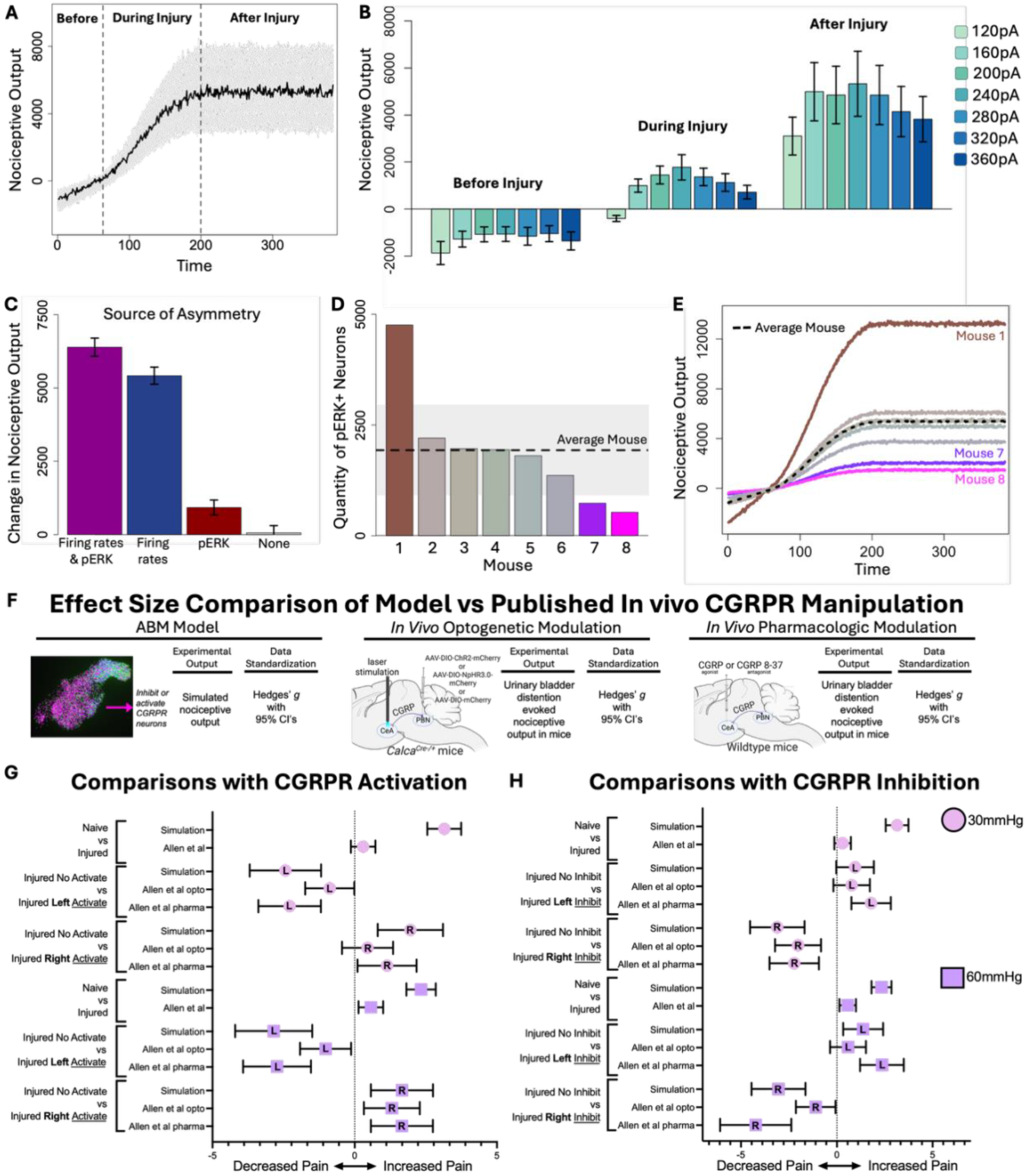
Model predictions of nociceptive output and comparisons with *in vivo* studies. **(A)** Simulated nociceptive output for the average mouse during constant 240 pA stimulation. The black line shows stochastic variation within a single simulation; gray shaded region represents variability across pERK expression levels (±1 mean absolute deviation, MAD). **(B)** Model predictions of nociceptive output for average mouse before, during, and after injury for constant noxious stimulations ranging from 120 pA to 360 pA. Nociceptive output increased significantly during and after injury across all stimulation intensities (p < 0.001), with largest relative increases occurring at low-to-mid stimulation levels (160 – 280 pA). Values represent mean ± 1 SD. **(C)** Predicted injury-induced increases in nociceptive output for models incorporating both firing rate and pERK asymmetries, firing rate asymmetry alone, pERK asymmetry alon±, and no hemispheric asymmetries. Values represent mean ± SD across 100 simulations. Simulations show firing rate asymmetry to be the main driver of changes in nociception due to injury. ± Quantity of pERK-expressing neurons for each mouse-specific model parameterization (n = 8 mice). The dashed line indicates pERK expression in the average mouse and gray shaded region shows ± 1 MAD. **(E)** Average nociceptive output from 100 simulations of each mouse-specific parameterization (n = 8). Nociceptive output was highest for parameterizations corresponding to mice with elevated pERK expression (e.g. Mouse 1) and lowest for parameterizations corresponding to mice with lower pERK expression (e.g., Mouse 7, 8). (**F**) In comparison experiments, the impact of manipulation of the CGRPR-expressing cells in the left or right CeA under control (naive) <u>or</u> CYP-sensitized (injured) conditions is evaluated on simulated model output (nociceptive output) or changes in system-level physiology responses during bladder distention from *Allen et al 2023*. In *Allen et al 2023*, both optogenetic and pharmacologic approaches were used to activate CGRPR-expressing cells or inhibit these cells. Optogenetics involved the expression of ChR2 or NpHR3.0 in cells that projected from the PBN to the CeA. Pharmacology involved delivery CGRP or CGRP 8-37 (antagonist) directly to the CeA. Data from dry and wet lab experiments were then converted to Hedges’ *g* standardized mean effects. (**G**) Forest plots show Hedges’ *g* 95% confidence intervals in wet-lab experiments when CGRPR-expressing cells are activated. Positive Hedges’ *g* is indicative of an increase in pain-like effect for the indicated comparison and a negative Hedges’ *g* is indicative of a decrease in pain-like effects. Pink data points are for low noxious stimulation (120 pA for simulation and 30 mmHg *in vivo*). Purple data points are for high noxious stimulation (240 pA for simulation and 60 mmHg *in vivo*). Top two comparisons for the pink and purple data show Hedges’ *g* calculations for no cellular manipulation experiment when CGRPR-expressing neurons are not manipulated. Simulated data shows a stronger impact of “injury” compared to mouse experiment at both low and high noxious stimulation. Remaining comparisons show indicated manipulation of the left or right CeA. (**H**) Forest plots show Hedges’ *g* ± 95% confidence intervals in experiments when CGRPR-expressing cells are inhibited. Simulated and experimental conditions are the same as in **G** but now CGRPR-expressing cells are inhibited with simulation, optogenetics, or pharmacology.

Model predictions of nociceptive output before, during, and after injury were compared across all stimulation intensities used in the wet-lab experiments (**Figure 4B**). There was a significant main effect of time (F(2,1240) = 8508.97, P<0.001), with nociceptive output increasing substantially from before injury to during injury and further increasing after injury. There was also a significant main effect of simulation intensity (F(6,1240) = 145.12, P<0.001), indicating higher stimulation intensities produced greater nociceptive output. Importantly, we observed a significant time x intensity interaction (F(12,1240) = 15.43, P<0.001) indicating that the effect of stimulation intensity was amplified during and after injury. Post hoc contrasts showed that this amplification was most pronounced at low-to-mid intensities (160–280 pA) and diminished at the highest intensities (320–360 pA). These data suggest a saturation of the injury-related increases in nociceptive output at high stimulation levels.

To determine which experimentally observed neuronal asymmetries most strongly influence nociceptive output, we simulated four model variants: asymmetry in both neuronal firing rates and pERK expression, asymmetry in firing rates alone, asymmetry in pERK expression alone, and a fully symmetric model with no hemispheric differences (**Figure 4C**). Hemispheric asymmetry in neuronal firing rates was the primary driver of injury-induced increases in nociceptive output. Equalizing firing rates across hemispheres substantially reduced nociceptive output, even when pERK expression remained asymmetric. In contrast, preserving firing rate asymmetry while equalizing pERK expression across hemispheres largely maintained injury-induced increases in nociceptive output. Together, these results indicate that hemispheric differences in neuronal excitability dominate CeA-level nociceptive signaling in the model. In the absence of any asymmetries, nociceptive output remained stable over time.

Finally, the number of pERK-expressing neurons varied considerably across animals (**Figure 4D**), which translated into a wide range of nociceptive predictions when individual mouse parameterizations were simulated (**Figure 4E**). Model configurations corresponding to animals with elevated pERK expression (e.g., Mouse 1) produced the greatest change in nociceptive output due to injury, whereas parameterizations corresponding to animals with lower pERK expression (e.g., Mouse 7 - 8) produced more modest responses.

### Standardized effect analyses demonstrate good concordance between model predicted effect sizes and observed effect sizes in mice

To validate the model, we tested whether model-predicted changes in nociceptive output during lateralized activation or inhibition of CGRPR-expressing neurons match our previously published *in vivo* findings from a CYP-induced bladder pain model in female mice (15). In the model, activation was implemented by setting neuron firing rates to 15 Hz, whereas inhibition was implemented by setting firing rates to 0 Hz. We simulated hemisphere-specific activation and inhibition at two stimulation intensities (120pA and 240pA), corresponding to low and high bladder distention pressures (30mmHg and 60mmHg) used in mice. Model simulations included all eight parameterizations (Mouse 1 – 8) to capture biological variability. Replicate model simulations were performed for each parametrization to align with sample sizes used in the *in vivo* experiments. Model output from before and after injury time points were evaluated to mirror the control and CYP-sensitized conditions *in vivo*. In the previous experimental studies, we used optogenetics and pharmacology to activate or inhibit CGRP/CGRPR-expressing cells in the left or right CeA in control (“intact”) and CYP-sensitized (“injured”) mice. Here, we directly compare *in silico* manipulations in the model with each *in vivo* approach (**Figure 4F**) using Hedges’ *g* mean effect size analysis to standardize outcomes across approaches.

We first examined the effects of targeted CGRPR-expressing neuronal activation. Effect sizes were quantified as standardized mean differences using Hedges’ *g* with 95% confidence intervals (CIs) and visualized using forest plots (**Figure 4G**). Values greater than zero are indicative of increased “pain.” We compared the standardized mean differences of injured and uninjured (naïve) pain output in the absence of additional activation. As expected, both wet-lab data and the model simulations yielded positive effect sizes at low (120pA/30mmHg) and high (240pA/60mmHg) stimulation levels. However, the magnitude of the pain effect from the model data was significantly higher than the in-vivo experiments. Next, we compared effect sizes between control and injured animals when left or right CGRPR-expressing cells were activated. Here, we found strong alignment between the model simulations and the pharmacology data at both stimulation intensities and for both the left and right-side manipulations. There was weaker agreement between the model and the optogenetic functional data; however, the CIs for all groups overlapped indicating no statistically significant differences.

Second, we ran an analogous comparison between the model and *in vivo* functional experiments with inhibition of the CeA CGRP/CGRPR-expressing cells (**Figure 4H**). Again, we found strong alignment between model simulations and the *in vivo* experimental data with the biggest outlier being the optogenetic experimental group, which tended to show the lowest overall effects. Overall, this standardized effect size comparison experiment demonstrated that predicted outputs from the bilateral ABM aligned well with experimental data.

## Discussion

Pain is shaped by interacting neuronal populations distributed across circuits that respond to sensory stimuli, produce emotional-affect, and appropriate behavioral changes. In this study, we integrate *ex vivo* physiology, spatial histology, and *in vivo* behavioral comparisons with a bilateral, three-dimensional agent-based model of the CeA to show that hemisphere-specific differences in intrinsic neuronal excitability, particularly within CGRPR- and PKCδ-expressing neuronal populations are dominant drivers of CeA nociceptive output in a bladder pain model. The model incorporates heterogenous wet-lab data and demonstrates that small biological differences can scale to large differences in pain behavior.

Prior studies indicate that the CeA can both amplify and suppress pain in a cell-type specific manner, with PKCδ-expressing neurons generally promoting nociception and SST neurons exerting antinociceptive effects (19). Subsequent studies showed hemisphere-specific nociceptive processing between the left and right CeA (15). Multiple molecular and physiological studies show a right-hemisphere dominance for pronociceptive processing, including stronger ERK and PKA pathway activation and larger injury-evoked increases in neuronal firing (7). By demonstrating that asymmetry in neuronal excitability outweighs other measured asymmetries, such as pERK expression, our model provides a systems-level explanation for the consistent behavioral impact of right-biased CeA activation across pain paradigms.

In visceral bladder pain, CeA output critically depends on mGluR5-linked plasticity (30) and strong parabrachial (PBN) input carrying glutamate and CGRP (15). CGRP signaling exerts opposing lateralized behavioral outputs where pain-like effects increase via the right CeA and decrease via the left CeA. This phenomenon is mirrored in our simulations. In inflammatory pain models such as formalin, CFA, or arthritis, CeA ERK1/2 activation drives long-lasting hypersensitivity, with right-biased ERK activity producing disproportionate pain output (12, 13). Electrophysiology data from arthritic models further shows stronger right-CeA sensitization and greater dependence on kinase signaling, supporting asymmetry in neuronal excitability as a principal determinant of pain output (7). In neuropathic pain, early injury effects rely on PBN to CeA synaptic plasticity and CRF projection neurons, whereas chronic stages shift toward hyperexcitability in non-CRF populations, suggesting that chronic pain represents a transition to a distinct emergent state (1, 31). Stress-related enhancement of CRF1 signaling in the CeA further strengthens excitatory BLA to CeA pathways (32) and intensifies pain, consistent with excitability-driven mechanisms (9). Interestingly, when directly comparing simulation predictions to wet-lab data, we found better concordance to pharmacologic studies compared to optogenetic experiments. This likely highlights a fundamental difference between such studies even if interpretation is often confabulated in the literature. Pharmacologic manipulation of a target provides a much more precise interrogation of a signaling pathway than optogenetics, and, in the context of computational design may be a better source of data for future iterations of the current model.

Our modeling framework makes several simplifying assumptions that define its scope. The ABM focuses on two well-characterized CeA cell populations, providing a foundation for integrating cell-type-specific physiology but not capturing the full molecular diversity of the CeA. Electrophysiology parameters are derived from *ex vivo* recordings in a single bladder pain model, which do not fully capture *in vivo* network influences. The ABM simplifies circuit architecture by assuming a reduced inhibitory connectivity structure and imposing fixed hemispheric valence, and it omits sex-specific parameters and descending pathways. In addition, pERK expression is used as a proxy of neuronal activation during noxious stimulation. Together, these assumptions provide a controlled framework for identifying dominant drivers of nociceptive output while establishing a foundation for future expansion to incorporate additional cell types and circuit interactions.

Our bilateral 3-D ABM advances prior amygdala modeling by incorporating realistic spatial organization and hemisphere-specific parameterization. By capturing animal-to-animal variability in firing rates, pERK counts, and cell distribution, the model identifies intrinsic excitability as the primary mechanism through which diverse molecular pathways ultimately modulate nociception. This is demonstrated by simulations showing that once intrinsic excitability is equalized across sides, hemispheric differences in nociceptive output collapse, revealing firing-rate asymmetry as the key driver despite persistent upstream signaling biases. Thus, our approach provides a compact systems-level hypothesis for the origin of CeA lateralization. Also, it provides a computational foundation for evaluation of the amygdala in other contexts such as fear conditioning and anxiety modulation, where known parameters can be used for simulations to derive the emergent factor that drives these behavioral outputs. There also exists potential for single models that integrate data across contexts. In other words, single computational approaches that include outputs for multiple contexts. That is aspirational but would help push all fields to intentionally consider overlap in cell populations and dependent variables in wet-lab experiments. Moreover, the ABM framework is flexible and can be adapted to essentially any neural area where there exists enough physiological and functional data to model a structure. Resources such as the Blue Brain Atlas (29) provide the relevant volumetric data necessary to complete this modeling in three dimensions, which is a critical parameter long ignored in many basic science studies of the rodent nervous system.

## Materials and Methods

### Animals (unpublished data)

We used adult (8–13 weeks) male and female *Calcrl*^Cre^ (Calcrl^tm1.1(cre)Rpa^; provided by Dr. Richard Palmiter, University of Washington (33), *Prkcd*^*Cre*^ (Tg(Prkcd-glc-1/CFP,-cre)EH124Gsat/Mmucd; provided by Dr. Yarimar Carrasquillo (34), and Ai14(RCL-tdT) reporter mice (Jackson Laboratory #007914). Parental lines were maintained as heterozygotes (*Calcrl*^Cre/wt^ and *Prkcd*^*Cre/wt*^). These mice were crossed with Ai14(RCL-tdT)-D^+/-^ animals (35) to generate F1 offspring, and heterozygous F1 mice (Calcrl-Cre; Ai14 (tdTomato) or Prkcd-Cre; Ai14 (tdTomato)) were selected for whole-cell patch-clamp recordings. In *ex vivo* slice experiments, CGRPR-expressing or PKCδ-expressing neurons were identified by TdTomato fluorescence. All lines were backcrossed to a C57BL/6J background. Mice were group-housed under a 12:12 light–dark cycle with food and water available *ad libitum*. Both males and females were included unless otherwise noted. All procedures were approved by the Institutional Animal Care and Use Committee (IACUC) at The University of Texas at Dallas (protocols 20-04 and 2023-0092) and conducted in accordance with institutional guidelines. All other animal research information used in the model was from published studies and can details can be found in the original studies.

### Cyclophosphamide Bladder Sensitization (unpublished data)

Cyclophosphamide (CYP; Sigma-Aldrich cat # C0768) is dissolved in sodium chloride 0.9% normal saline and was used to induce bladder pain-like sensitivity in mice. Mice received 100 mg/kg CYP (intraperitoneally) every other day for 5 consecutive days (3 total injections). 24 hours after the first injection of CYP, is considered 1-day post-injury (1 DPI). The experimenters were blinded to the treatment.

### Ex vivo electrophysiology (unpublished data)

#### Acute slice preparation

After 3 doses (across 5 days) of CYP treatment or saline control, at 6/7 DPI, Calcrl-Cre; Ai14 (tdTomato) or Prkcd-Cre; Ai14 (tdTomato) mice were decapitated and brains were rapidly extracted, placed in ice-cold cutting solution, and cut in coronal slices (250–300 μm) using a Leica VT1200 S vibrating blade microtome (Leica Microsystems Inc.). The cutting solution was composed of the following: 110 mM choline chloride, 25 mM NaHCO_3_, 1.25 mM NaH_2_PO_4_, 2.5 mM KCl, 0.5 mM CaCl_2_, 7.2 mM MgCl_2_, 25 mM D-glucose, 12.7 mM L-ascorbic acid, and 3.1 mM pyruvic acid, oxygenated with 95%/5% O_2_/CO_2_. The slices containing CeA were incubated at 25°C for at least 60 min in a holding chamber containing artificial CSF (ACSF) composed of the following: 125 mM NaCl, 2.5 mM KCl, 1.25 mM NaH_2_PO_4_, 25 mM NaHCO_3_, 2 mM CaCl_2_, 1 mM MgCl_2_, and 25 mM D-glucose. The slices were then moved to the microscope bath and recovered for at least 10 min at 33°C before recording. During incubation and recovery, the chambers were continuously oxygenated with 95%/5% O_2_/CO_2_.

#### Whole-cell patch-clamp recordings

The recording chamber was perfused continuously with ACSF oxygenated with 95%/5% O_2_/CO_2_ (1 ml/min) and all recordings were performed at 33 ± 1°C. A recording chamber heater and an in-line solution heater (Warner Instruments) were used to control and monitor the bath temperature throughout the experiment. Recording pipettes (3.5- to 6.6-MΩ resistance) were filled with internal solution composed of the following: 120 mM potassium methyl sulfate, 20 mM KCl, 10 mM HEPES, 0.2 mM EGTA, 8 mM NaCl_2_, 4 mM Mg-ATP, 0.3 mM Tris-GTP, and 14 mM phosphocreatine with pH 7.3 using 5 M KOH and an osmolarity of ∼300 mosmol^−1^. Whole-cell current-clamp recordings were obtained from tdTomato-expressing CeA neurons located in the capsular division (CeC) or lateral division (CeL) in the right or left hemisphere. Cells were visually identified using an upright microscope (Leica DM6 F6) equipped with differential interference contrast optics with infrared illumination and epifluorescence. Recording electrodes were visually positioned in the CeC, guided by the distinctive fiber bundles and anatomic landmarks delineating its structure. Recordings were controlled using the Multiclamp 700B patch-clamp amplifier interfaced with a Digidata 1440A acquisition system and pCLAMP 10.7 software (Molecular Devices) on a Dell computer. Before forming a membrane-pipette seal, pipette tip potentials were zeroed. Whole-cell capacitance was derived offline from membrane time-constant calculation obtained from capacitance curve traces in current-clamp configuration. Spontaneously active cells were recorded gap-free in current-clamp configuration. Position of cells ranged from -1.22 to -1.70 mm of Bregma. Recordings were completed blinded to treatment condition. 500 ms long depolarizing current of various amplitudes were injected from resting membrane potential to cells that were silent at rest, to elicit repetitive action potential firing. Liquid junction potentials were not corrected for the recordings. All recordings were acquired at 20 kHz and filtered at 4 kHz. Recording sites were constructed using the mouse brain atlas as a guide (<u>Paxinos et al., 2001</u>). Position of cells ranged from -1.22 to -1.70 mm of Bregma.

#### Electrophysiology Data Analysis and Statistics (unpublished data)

Minimum sample sizes used in each experiment were based on previous studies of the CeA in nociception. Cells were allocated into experimental groups based on saline-treated and CYP-treated groups, which were further categorized into left and right CeA. Electrophysiological data were analyzed using ClampFit 11.4 (Molecular Devices), Microsoft Excel, Mini Analysis (v. 6.0.8, Synaptosoft), and Prism (version 10.4.1, GraphPad Software Inc.). Neurons with late-firing phenotype were identified by the sharp rising edge followed by a delay in firing when 500 ms depolarization current steps were injected. Neurons of regular-spiking phenotype were identified by the smooth rising phase with no delay in firing when 500 ms depolarization current steps of increasing intensity was injected. Another distinguishing factor to identify late firing vs. the regular spiking phenotype along with the sharp rising edge is the higher rheobase current required by the late firing neurons. The spontaneously firing phenotypes did not need any current injection. Adaptation of inter spike interval (ISI) was calculated from the ratio of the measurements obtained from the last and first action potential in response to a 500-ms depolarizing current injection at 2× rheobase. ISI accommodating cells were defined as cells with a ratio ≥1.5, whereas ISI non-accommodating cells had a ratio of <1.5. Firing frequency was determined as the number of spikes evoked when 500 ms depolarization current steps of incremental increase of 40 pA from 0 pA to 360 pA was injected into the neuron. The methods for obtaining resting potential, capacitance, rheobase, spike amplitude, rise time, spike duration, current threshold (I-Threshold), voltage threshold (V-Threshold) are provided in the **Supporting Information**.

Results are expressed as mean ± SEM. For results that have one dependent variable and two groups, t tests were chosen based on their variance (Welch’s correction if unequal variance) and normal distribution (unpaired t test if normally distributed, or Mann-Whitney U test’s if non-normally distributed). For the results that have two independent variables and two groups, two-way ANOVA followed by post-hoc Tukey’s multiple comparison test was used. Fisher’s exact test was used to compare the proportion of distribution of neuronal sub-types. p values<0.05 were considered significant. All statistical analyses were performed using GraphPad Prism (version 10).

### Western Blot Analysis (unpublished data)

Wildtype female C57Bl/6J mice were treated with CYP (or control saline) as described above. On 6 DPI (1 day after the last CYP treatment), mice were given urinary bladder distention (UBD) using our previously published protocols(6). Briefly, following urethral catheter insertion under full isoflurane anesthesia, isoflurane was reduced over 45 min to a low level where animals responded to toe pinch. UBD was then performed with an escalating set of bladder distention pressures in the following order: 15, 30, 45, 60, 75 mmHg. The bladder was distended for 20 sec at each pressure in triplicate with a 2 min intertrial interval. Total experimental time during distention trials was 35 min. 5 min following distensions, animals were decapitated, brains were removed, sectioned into 1 mm thick coronal sections using an acrylic brain matrix (Stoelting), and 1 mm diameter amygdala punches were obtained from both sides of the brain using a punch tool (Stoelting) then stored on dry ice. Samples were homogenized with ice-cold homogenization buffer (20 mM Tris-HCl, pH 7.5, 1 mM EDTA, 1 mM Na4P2O7, 100 μM PMSF, 25 μg/mL aprotinin, 25 μg/mL leupeptin, Phosphatase Inhibitors II and III (Sigma Aldrich Cat P5726 P0044) then assessed for total protein content using the Pierce BCA protein assay kit (Thermo Scientific Cat A55860). 10 μg of each sample was separated on a 12.5% NuPAGE gel (Thermo Scientific Cat WBT41212BOX) then transferred to nitrocellulose membrane. Membranes were blocked in Odyssey blocking buffer (Licor Cat 927-70001) for 1 hr then co-incubated with mouse anti-pERK1/2 (1:1,000, Cell Signaling Cat: 9106L) and rabbit anti-ERK1/2 primary antibodies (1:1,000, Cell Signaling Cat: 9102L; in 0.1% Tween 20 in Odyssey blocking buffer) for 1 hr. Blots were rinsed with 0.1% Tween 20 in TBS (TTBS) then incubated with goat anti-mouse Alexa 680 (1:20,000, Molecular Probes Cat: A-21058) and goat-anti rabbit IR 800 secondary antibodies (1:20,000, Rockland Cat: 11-132-002; in 0.1% Tween 20 in Odyssey blocking buffer) for 1 hr. Following a final rinse in TTBS, blots were scanned on an Odyssey infrared imaging system. pERK1/2 band densitometry in each sample was normalized to total ERK1/2 using Image Studio Lite software (version 4, LI-COR). Quantification was done in a blinded fashion.

Saline-treated and CYP-treated groups were independently evaluated with two-way ANOVA (hemisphere side and distention condition as independent variables) followed by post-hoc Bonferroni multiple comparisons. p values<0.05 were considered significant. Statistical analyses were performed using GraphPad Prism (version 10).

### Cell Dendrite Length Calculations (published data)

Previously, we were able to establish the importance of proper connectivity within the model, as it runs primarily on a firing-rate equation that is based on the location of the cells. The location of the cells can be further specified with dendrite endpoint distance data, which helps distinguish between the cell body and the dendrite. In our previous publication (26), we evaluated the overall morphology of PKCδ-expressing neurons using biocytin cell filling after patch clamp recording. Briefly, cells were filled and the slice was fixed in 4% PFA. Samples were then incubated overnight, at 4°C and protected from light, in 1:500 Alexa Fluor 647 streptavidin (The Jackson Laboratory 016-600-084) in blocking solution containing 1.5% normal goat serum (NGS; Vector Labs), 0.1% Triton X-100, 0.05% Tween 20, and 1% bovine serum albumin (BSA). In minimal light, slices were then washed in 0.1 m PBS four times for 30 min at room temperature. Slices were then cleared using increasing concentrations of 2,2’-thiodiethanol (TDE) for 10 min each using 10%, 30%, 60%, and 80% concentrations, followed by incubation in 97% TDE for 2 h. Slices were then mounted on positively-charged glass slides and covered with glass coverslips using 97% TDE.

Images of recovered biocytin-filled neurons were taken using a Nikon A1R laser scanning confocal microscope with a 40x oil-immersion objective. Gain and pinhole size were kept constant between experiments. Sequential acquisition of z-stacks was collected at 0.09-μm steps. Images were collected at varying sizes, depending on the extension of the dendrites of each neuron, and were then automatically stitched on acquisition using NIS Elements software. The images were then automatically stitched using NIS Elements software and analyzed on NeuroLucida 360 (MBF Bioscience). In the present set of experiments, we reanalyzed the total lengths of all processes from the cells.

The NIS files were opened on Blender (Blender Foundation www.Blender.org Version 4.2.1 LTS) for 3-D visualization and then labeled and measured. Each file had one PKCδ-expressing cell with different dendrites and dendrite endpoints. The dendrite endpoints were first labeled for better identification. As an example, in **Figure 3G, 3H**, the endpoints are labeled based on length, with 1 being the longest dendrite and 3 being the shortest. Then, the subbranches are also labeled based on length. For example, in **Figure 3G**, branch 1 is then split into branch 1A and 1B, with 1A having the longest endpoint. 1A is once again divided into 1A1 and 1A2, based on the length of the dendrite **(Figure 3I)**. This format was continued until all the endpoints were labeled within the Blender file and continued for all the PKCδ-expressing cells originally filled in our previous publication. For measurements, a straight line was drawn from the center of the cell body to the end of the dendrite. The measurement was taken by rotating the model in all directions to ensure the correct length was taken, considering the 3D space. Once the measurements were taken, they were recorded based on each PKCδ-expressing cell in each file at each 10μm interval from the radius.

### Whole animal physiology evaluation of CGRPR-expressing neurons in the CeA in a model of bladder injury (published data)

Finally, we utilized our previously published behavioral data showing the impact of manipulating CGRPR-expressing neurons in control and CYP-sensitized injured female mice (15). Following CYP treatment, mice were tested for mechanical sensitivity and/or put through the urinary-bladder distention visceromotor response (UBD-VMR) assay. Most relevant to the present manuscript, CYP-treated mice exhibit mechanical hypersensitivity and increased UBD-VMRs. To manipulate CGRPR-expressing neurons, we utilized inhibitory and excitatory optogenetic vectors. Viral-based cre-recombinase dependent vectors (rAAV5/Ef1a-DIO-hCHR2-mCherry, rAAV5/EF1a-DIO-NpHR3.0-mCherry, or rAAV5/EF1a-DIO-mCherry) were injected into the right or left CeA of female mice from the *Calca*^*Cre/+*^ *(Calca*^*tm1*.*1(cre/EGFP)Rpa*^) transgenic strain (15, 36). As described in the introduction above and previously reported, we found that CGRP was largely anti-nociceptive in the left CeA and pronociceptive in the right CeA. This corresponded to Figures 1E-F, 1H-I, 1K-L and 1N-O from Allen et al, 2023 (15). These data were used to validate system-level output generated from our ABM. For comparisons between experimental data and model output, we utilized a Hedges’ *g* effect size calculation (see *Standardized mean effect analysis* section below).

### 3D Agent-Based Model of CeA Neurons

We developed a 3D agent-based model (ABM) of the left and right mouse CeA to determine how hemisphere-specific changes in neuronal excitability and pERK expression of PKCδ-expressing and CGRPR-expressing neurons during bladder injury influence nociceptive signaling. The model was implemented in NetLogo3D (version 6.4.0) (37) and is publicly accessible through our web application (23). All source code and input files are publicly accessible on the Open Science Framework (doi: 10.17605/OSF.IO/SWRK8). A comprehensive description of the model, including all assumptions and parameter values can be found in the **Supplemental Information**.

The model’s spatial domain is a 3-D reconstruction of the mouse CeA and its subnuclei derived from the Blue Brain Cell Atlas (29). Each CeA is initialized with approximately 10,000–13,000 agents representing individual PKCδ- or CGRPR-expressing neurons. All neurons are assigned variables tracking individual properties (**Table S1**) while global variables track system-level properties (**Table S2**). Neurons are assigned locations, cell-type expression identities (PKCδ, CGRPR, or both), firing types (regular spiking, late firing, or spontaneous), and initial pERK expression levels based on experimental distributions (**Figure S5**). An inhibitory network of connections between neurons is established using a stochastic algorithm, with connection probabilities derived from empirically observed log-normal distributions of dendrite lengths (**Figure S4**). Noxious stimulation is represented in the model as identical current injection (pA) to neurons in both the left and right CeA.

During each simulation time step, neuron-level properties are updated to reproduce the hemisphere-specific injury changes observed experimentally in PKCδ-expressing and CGRPR-expressing populations. Neurons accumulate sensitization through a damage variable (0 ≤ *d* ≤ 100) that increases during periods of noxious stimulation (≥120 pA). As damage increases, neuronal firing rates transition in response to injury according to hemisphere-specific distributions estimated from control and CYP-injured mice (**Table S3**). Similarly, additional neurons are recruited to express pERK in response to bladder distention and injury in a hemisphere-dependent manner, capturing the observed left-right differences in pERK expression (**Figure S6**). Neurons send inhibitory signals to one another through the connectivity network. Individual neurons are inhibited (i.e., firing rate set to 0 Hz) when their total inhibitory input exceeds a fixed threshold (15 Hz).

Emergent nociceptive output is computed each time step by aggregating firing rate activity across all pERK-expressing neurons. In the left CeA, PKCδ-expressing and CGRPR-expressing neurons are assumed to exert antinociceptive effects, whereas in the right CeA the same populations are assumed to exert pronociceptive effects, reflecting opposing functional roles across hemispheres (15, 26, 28). At each time step, contributions to nociceptive signaling from the left and right CeA, respectively, are calculated as

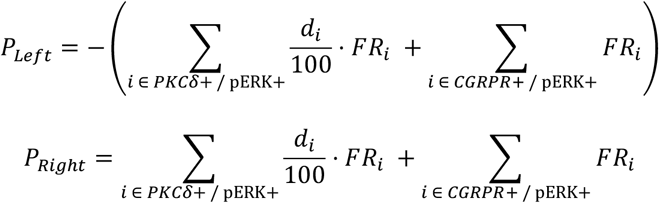

where *FR*_*i*_ denotes the firing rate of neuron *i* and *d*_*i*_ represents its sensitization (damage) state. As in our prior models, the contributions of PKCδ-expressing neurons are scaled by damage. Total nociceptive output at each time step is defined as *P*_*Total*_ = *P*_*Left*_ + *P*_*Right*_. This formulation produces a system-level measure of nociceptive output that emerges from noxious stimulation, hemispheric differences in firing rates and pERK expression, and inhibitory network dynamics.

Model stochasticity arises from random initialization of neuron-level parameters and from firing rates sampled from Poisson distributions with means calibrated to experimental electrophysiological recordings. To account for this variability, each simulation was replicated 50-100 times, and summary statistics were computed from the distribution of emergent outcomes. All statistical analyses were performed using R statistical software (38).

### Standardized Mean Effect Analysis

Standardized mean effect size analysis was completed to directly compare predicted outcomes from the bilateral ABM and our wet-lab experiments. For our wet-lab experiments, we used data from published behavioral experiments designed to look at the impact of CGRPR-expressing neuronal inhibition or excitation with inhibitory or excitatory optogenetics or pharmacology on mechanical sensitivity in control or CYP-treated mice (15). We replicated the conditions of these wet-lab experiments using our ABM by silencing (or activating) the appropriate CGRPR-expressing neurons in the model and recording nociceptive output at times t = 50 (before simulated “damage accumulation”) and t = 300 (after “damage accumulation”) from 8 simulations across 8 mouse parameterizations. We chose to use 8 total model simulations to match the number of animals used for PKCδ and CGRPR expression data and because the average sample size (n) in the wet-lab experiments was 8.176 with a standard deviation of 1.740. We also compared the no manipulation “intact” model (no inhibition or activation) to treatment control or CYP-treated mice without manipulation. These wet-lab data had an average sample size (n) of 48 so wet lab data were compared to six replications of each the eight mouse parameterizations (48 total simulations) in the ABM.

In all simulations, we assumed a constant 120 pA or 240 pA current corresponding to bladder distention pressure values of 30mmHg or 60mmHg in wet-lab experiments, respectively (see **Table S5** for results of these simulations and corresponding *in vivo* data). The y-axis is different in our wet-lab experiments (visceral-motor response) compared to the model output (arbitrary “Total Nociceptive Output” units). To normalize these axes, we calculated standardized mean effect sizes. Standardized mean effect sizes were calculated using a Hedges’ *g* value and 95% confidence intervals allowing for comparison between model outputs with constant number of replicates (n = 8 for inhibition/activation and n=48 for intact comparisons) and wet-lab data with a variable number of samples per group (see **Table S5** for Hedges’ *g* data). The Hedges’ *g* value was calculated as

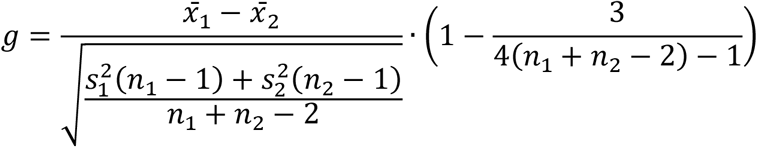

where 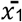 is the treatment group mean and 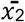 is the control group mean, s_1_ and s_2_ are respective standard deviations, and n_1_ and n_2_ are respective sample sizes. All calculations were done using Cambell Collaboration’s Effect Size Calculator (32).

## Acknowledgments

We acknowledge funding from the National Institutes of Health (NIH) including R01 DK115478 (BJK/RMN), R15 NS128624 (RMN/BJK), and the Intramural Research Program of the National Center for Complementary and Integrative Health, NIH (YC) along with additional support from the University of Texas at Dallas URAP program (CN/BJK) and the Duquesne University Neurogenerative Undergraduate Research Experience (NURE) program (MK/FZ) (R25 NS100118).

