## Supporting Information for "Predicting Pain Using a Data-Driven Agent-Based Model of the Bilateral Central Amygdala"

**This PDF file includes:**

Supporting Text: Model Description
Figures S1 to S6
Tables S1 to S5
SI References

### Supporting Information Text

#### Supplementary neuronal excitability measurements method description

Resting potential was measured from the initial section of the voltage waveform before injecting the depolarizing current. Rheobase is defined as the minimum current required to induce an action potential in response to a 500-ms depolarizing current injection for both late-firing and regular-spiking neurons. Whole-cell membrane capacitance was calculated by curve-fitting the voltage traces from the capacitive transients elicited by ±20 current steps. The voltage transient's exponential phase was used to derive the membrane time constant by a fitting method utilized by Levenberg-Marquardt algorithm in Clampfit software. Input resistance ($R_{in}$) was calculated using the average change in membrane potential in response to a ±20pA current injection of 500-ms duration. Whole-cell capacitance is then calculated from the following equation:

$$T_m = R_m{*}C_m,$$

where $T_m$ = membrane time constant, $R_m$ = input resistance, $C_m$ = membrane capacitance.
Single action potential properties were measured from the action potentials generated in response to a 5-ms depolarizing current injection in the current-clamp mode. Action potential amplitude was measured from the threshold potential to peak of the rising phase. Rise time of the action potential was defined as the time required for the membrane potential to reach peak voltage from threshold potential. Action potential duration (APD) was measured at 100% repolarization to threshold potential. Current threshold for action potential generation was defined as the minimum current injection required to elicit the action potential. Voltage threshold of the single action potential was measured manually from the traces **(Figure S1, S2)**.

#### ABM Model description

Below is a detailed description of our 3-D agent-based model (ABM) of PKC$\delta$- and CGRPR-expressing neurons in the left and right central nucleus of the amygdala (CeA). This section is written in accordance with the Overview, Design concepts, and Details (ODD) protocol (Grimm et al., 2020), which is a standard format for describing ABMs. The model is publicly accessible through our web application (Perumal et al., 2025). All source code and input files are available on the Open Science Framework (doi: 10.17605/OSF.IO/SWRK8).

**1. Purpose of Model.** The purpose of the ABM is to investigate how hemisphere-specific changes in CeA neuron excitability and pERK expression due to bladder injury influence pain signaling from the CeA. The ABM was developed in NetLogo3D (Version 6.4, Wilensky, 1999). The model simulates the connectivity, firing rates, and pERK expression of PKCδ- and CGRPR-expressing CeA neurons in the left and right CeA and produces an emergent measure of nociceptive output from the bilateral CeA. Model parameters were estimated from experimental data demonstrating left-right hemisphere differences in CeA neuron firing rates and pERK expression. Model simulations were performed to quantify how these hemisphere-specific differences contribute to pain signaling at the CeA level.

**2. Entities, state variables, and scale.** The model's spatial domain is a 3-D reconstruction of the mouse CeA derived from the Blue Brain Cell Atlas (Erö et al., 2018). Mesh files (.obj) defining the surface coordinates of the right mouse CeA and its subnuclei were obtained from the Blue Brain Cell Atlas and rendered using Blender software (version 4.2 LTS, Blender Foundation, 2024). The interior of each subnuclei was sampled in Blender and the resulting values were imported into NetLogo3D to determine the patches corresponding to the CeA and its subnuclei. The spatial domain for the right CeA in NetLogo3D consists of 32,412 patches (i.e., small cubes), each representing 25 $\mu$m x 25 $\mu$m x 25 $\mu$m. Patch colors denote the subnuclei: red for Capsular (CeC), blue for Lateral (CeL), and green for Medial (CeM) (**Figure S5**). The left CeA is represented in the model as a mirror image of the right CeA.

Each agent in the model represents an individual neuron in the CeA that expresses protein kinase C δ (PKCδ), calcitonin gene related peptide receptor (CGRPR), or both. The model also tracks which of these neurons express a phosphorylated extracellular signal releasing kinase (pERK). Each agent (i.e., neuron) has sixteen variables (**Table S1**) defining its properties and behaviors.

During initialization, each neuron is assigned a 3-D location (*xcor*, *ycor*, *zcor*) within the CeA and binary variables (*PKC?, CGRPR?, pERK?)* classifying its expression type. Expression of PKCδ and CGRPR are assigned at initialization and do not change during simulation. A subset of neurons express pERK at initialization and additional neurons are recruited to express pERK during simulated periods of distention and injury. Each neuron is assigned a firing rate type (*Rate-Type*) equal to regular spiking (RS), late firing (LF), or spontaneous (Spont). Neuron firing rate types are assigned at initialization and may change during the simulation. Each neuron has three variables related to its "damage." The damage variable (*d*) track's the neuron's sensitization caused by noxious stimulation. The rate at which a neuron accumulates damage during noxious stimulation depends on its latency period (*Latency*) and sensitivity (*Sensitivity*).

Each neuron also has five variables related to its connectivity with other neurons within the CeA. Two of these variables are the number of incoming connections (*Incoming-Connections*) and number of outgoing connections (*Outgoing-Connections*) for the neuron. In addition, the variables *Incoming-Connections-IDs* and *Outgoing-Connections-IDs* store a list of agent IDs associated with the neuron's incoming and outgoing connections, respectively. The fifth variable is the neuron's inhibition status (*Inhibited?*), which is a binary variable indicating whether the neuron is inhibited or not. Lastly, each neuron has a firing rate (*FR*) describing the frequency in hertz (spikes per second) of its action potential. Neuron firing rates are updated each time step.

Additionally, there are six global variables (**Table S2**). Global variables include the noxious stimulation intensity (*Stim*), total time under stimulation ($S_{Total}$), nociceptive output from the left CeA ($P_{Left}$), nociceptive output from the right CeA ($P_{Right}$), and total nociceptive output ($P_{Total}$). Additionally, a global variable (*Avg-Damage*) tracks the average damage across all neurons. Each model tick represents one time step.

**3. Process overview and scheduling.** The user must select several parameters via buttons on the user interface prior to model initialization. Specifically, the user must identify the CeA hemisphere (Left or Right), the spatial distribution of neurons (corresponding to Mouse ID 1 through 8 or Average Mouse), and the intensity of noxious stimulation (120 pA, 160 pA, 200 pA, 240 pA, 280 pA, 320 pA, or 360 pA). After these parameters are selected, the model can be initialized and simulated.

Initialization (SETUP procedures)
The following procedures are executed once during initialization:

I. *Set-Up-Amygdala:* Import patch coordinates associated with CeC, CeL, and CeM and create spatial domain representing the CeA.
II. *Setup-Turtles:* Import spatial locations of PKCδ and CGRPR neurons for the selected mouse ID and create agents representing these neurons.
*III.* *Initial-pERK:* Identify initial subset of PKCδ and CGRPR neurons that express pERK.
*IV.* *Setup-Rate-Types:* Assign *Rate-Type* values to all neurons and create agentsets based on expression type and firing frequency (e.g., PKC-RS-neurons).
*V.* *Setup-Turtles-Damage:* Assign initial values of damage-related variables to all neurons.
VI. *Create-Network:* Create a network of connections between neurons using stochastic algorithm.
VII. *Load-Data:* Import mean firing rate data from Control and CYP-injured mice (**Table S3**) and create lookup tables.
*VIII.* *File-Opening-Setup:* Read in stimulation intensity file.

Simulation (GO procedures)
The following procedures happen each time step in the order listed below:

I. *Update-Damage:* Update damage variable (d) for each neuron.
*II.* *Update-pERK:* Select subset of non-pERK neurons to express pERK during distention (pre-injury) and injury phases.
*III.* *Update-Rate-Types:* Change *Rate-Type* of select neurons during injury phase.
*IV.* *Assing-Firing-Rates:* Update the firing rate variable (*FR*) of all neurons using stochastic process with mean firing rates stored in tables.

*V.* *Send-Inhibitory-Signals:* Send inhibitory signals through the network and update receiving neuron firing rates accordingly.
*VI.* *Calculate-Pain:* Compute emergent nociceptive output from CeA.
*VII.* Increment tick counter and update interface displays.

**4. Design Concepts**

**4.1 Emergence.** The primary emergent property of the model is the total bilateral nociceptive output, which evolves over time and in response to noxious stimulation. Nociceptive output is calculated separately for the left and right CeA and then summed for the total nociceptive output. Nociceptive output is calculated at each time step from neuron firing rates (Section 6.6).

Other emergent features of the model include the size of the connectivity network (i.e., the total number of connections between neurons), the average number of incoming and outgoing connections for PKCδ and CGRPR neurons, respectively, and overall inhibition of neurons through the network.

**4.2 Adaptation.** Neuronal adaptation is represented in the model through a damage accumulation process, which allows neurons to transition from an unsensitized (control) to a sensitized (injured) state over time when exposed to sustained noxious stimulation (≥120 pA). All neurons begin with zero damage ($d$ = 0) and accumulate damage at during periods of bladder distention (Section 6.2). As damage increases, neurons progressively shift toward a sensitized state, and their firing rates and rate types are adjusted accordingly. A neuron is considered fully sensitized when its damage variable reaches the maximum threshold ($d$ = 100).

Adaptation is further captured through changes in pERK expression. During both the distention (pre-injury) and injury phases, a subset of neurons that do not initially express pERK are recruited to express pERK (Section 6.3). These increases in pERK expression within the CeA vary by hemisphere and reflect experimentally observed adaptations to sustained bladder distention and injury.

**4.3 Interaction.** Neuronal interaction in the model occurs through inhibitory signaling. During each time step, a neuron may receive inhibitory inputs from one or more other neurons. If the sum of these inhibitory signals exceeds the inhibition threshold, the target neuron is suppressed for that time step (i.e., $FR$ = 0). The inhibition threshold is fixed at 15 Hz in all simulations. Neurons can transmit inhibitory signals only to other neurons within the same hemisphere. No cross-hemisphere neuron-to-neuron interactions occur in the model.

**4.4 Stochasticity.** Stochasticity occurs at multiple time points during the model's initialization and simulation procedures. During initialization, neurons are randomly assigned an (*xcor, ycor, zcor*) location within a specified cross-sectional slice of the CeA (Section 5). Values of damage sensitivity and latency parameters are randomly determined for each neuron using a uniform probability distribution with ranges displayed in **Table S1**. The connectivity network is established using a stochastic algorithm that creates connections between CeA neurons based on their distance from one another (Section 6.1). During each time step, the firing rates of PKCδ and CGRPR neurons are stochastically updated using values drawn from Poisson distributions with means calculated from experimental data (Section 6.4).

To account for the model's stochasticity, simulations are repeated 50-100 times, and the mean and standard deviation are reported for all emergent measures.

**5. Initialization.** The model is initialized by creating the CeA spatial domain and generating agents that represent individual neurons. The spatial domain is defined by reading values from three text files (.txt) that contain the x, y, and z coordinates of the patches within each of the three CeA subnuclei (CeC, CeL, CeM). Patches within each subnuclei are assigned the same color (red = CeC, blue = CeL, green = CeM). The entire CeA consists of 32,412 patches, each representing 25 $\mu$m x 25 $\mu$m x 25 $\mu$m. The overlay showing the CeA patches can be toggled off via the interface to display the neurons (**Figure S5**).

Depending on the mouse ID selected, the model generates approximately 10,000 to 13,000 agents, each representing an individual neuron. Neuron locations are imported from a text file, which specifies the number of PKCδ, CGRPR, and pERK expressing neurons in each cross-sectional slice of the CeA along the anterior to posterior axis (x = 4 to x = 70) (**Figure 3A-C**). For each slice, the appropriate number of neurons are created and assigned expression values (*PKC?* = yes or no*, CGRPR?* = yes or no) as well as a location within the corresponding CeA cross-section. Neurons are assigned a random location within the assigned CeA cross-section, with 80 - 90% of neurons assigned to patches in the CeL/CeC sub-nuclei and the remaining neurons assigned to patches in the CeM. Neurons are assigned a color based on expression type (PKCδ = blue, CGRPR = red, PKCδ and CGRPR = green) (**Figure S5**). Each neuron is also labeled to indicate whether it expresses pERK (*pERK?* = yes or no). The quantity of neurons that initially express pERK varies by Mouse ID (**Figure 3D-F**) and hemisphere (**Figure S6**).

Next, the firing rate type (*Rate-Type* = LF, RS, or Spont) of each neuron is assigned. The distribution of firing rate types differs between the PKC$\delta$ and CGRPR populations as well as by hemisphere. The initial percentage of neurons with each firing rate type is determined by experimental data from uninjured (control) animals (**Figure 1E, F**, **Figure 2E, F**). For neurons that co-express PKCδ and CGRPR, half are assigned PKCδ attributes and the other half assigned CGRPR attributes. Agentsets are created for each expression and rate type combination (e.g., *All-CGRPR-LF-turtles*, *All-PKC$\delta$-Spont-turtles*). Neurons co-expressing PKCδ and CGRPR are included in the appropriate agentset according to their assigned properties.

Finally, each neuron is initialized with zero damage (*d* = 0). Values for the *Latency and Sensitivity* variables are drawn from uniform distributions with ranges specified in **Table S1**.

After all neurons are initialized, connectivity is established using a stochastic algorithm (Section 6.1). The maximum number of connections per neuron can be adjusted via a slider on the interface. In all simulations, each neuron was limited to a single connection with another PKC$\delta$ or CGRPR neuron in the same hemisphere.

Lastly, multiple input files are read into the model, and the data are organized by type. Firing rate averages are stored in tables, while the intensity of noxious stimulation at each time step is stored as a list.

### 6. Submodels

**6.1 Create connectivity network (Create-Network).** Neuronal connectivity is established using a stochastic, distance-dependent algorithm. Each neuron begins with two empty lists, *Incoming-Connections-IDs* and *Outgoing-Connections-IDs*, which are updated as connections are formed. The likelihood of a connection between two neurons depends on the Euclidean distance (i.e. straight-line distance) between the neurons' centers. Data from reconstructed PKCδ neurons shows that connection lengths follow a log-normal distribution (**Figure S4**). Based on this distribution, we estimated the connectivity probabilities for four distance ranges: short (<30 μm, ~8%), medium (30–120 μm, ~50%), long (120–200 μm, ~27%), and extra-long (200–270 μm, ~14%). During the creation of the network, each neuron searches for neighboring neurons and randomly selects a neighbor to form a connection with based on these probabilities. When a connection is formed, the connecting neuron records the target neuron in its outgoing ID list, and the target records the connecting neuron in its incoming ID list.

**6.2 Update neuron damage (Update-Damage).** At initialization, the damage variable (*d*) for every neuron is set to 0, indicating the neuron is unsensitized. This is the equivalent of a control animal before injury. A neuron accrues damage only when the cumulative amount of time under stimulation exceeds the neuron's latency period ($S_{tot} > Latency$) and the stimulation intensity (*Stim*) is greater than or equal to 120 pA. Damage stops accumulating when it reaches its maximum value ($d = 100$), indicating the neuron is fully sensitized. This is the equivalent of an injured animal. For each neuron, damage at the time step $t$ is updated as:

$$d = \begin{cases} \min\left(d_{t-1} + \dfrac{100}{Sensitivity}, 100\right) & \text{if } S_{tot} > Latency \text{ and } Stim \geq 120 \\ d_{t-1} & \text{if } S_{tot} \leq Latency \text{ or } Stim < 120 \end{cases}$$

where $d_{t-1}$ is the value of the damage variable at the previous time step, $Sensitivity$ is the length of the neuron's sensitization period, and $Latency$ is the length of the neuron's latency period.

**6.3 Recruitment of pERK expressing neurons (Update-pERK).** The percent of neurons that express pERK (% number of neurons with pERK? = yes) increases during both bladder distention (pre-injury) and injury. The rate of increase in pERK expression varies by hemisphere but is the same across all mice.

During the pre-injury distention phase (yellow shaded region in **Figure S4**), neurons are selected at random to transition to a pERK-expressing state (pERK? = yes). Over this time period, pERK expression increases linearly, with an 18% increase in the left hemisphere and a 41% increase in the right hemisphere.

During the injury phase (orange shaded region in **Figure S4**), additional neurons are recruited at random to express pERK. In this phase, pERK expression increases by 26% in the left hemisphere and by 17% in the right hemisphere. Unlike pre-injury distention, increases in pERK expression during injury are proportional to average damage.

**6.4 Update neuron firing frequencies (Update-Rate-Types).** Neuron firing rate types (RS, LF or Spont) are updated every time step during periods of injury. Changes in rate types occur throughout the injury period in a manner similar to pERK recruitment. The percent of neurons in each rate type varies by cell-type and hemisphere but is the same across all mice.

In the right hemisphere, the PKCδ population is initialized with 48.5% RS, 31.4% LF and 20% Spont. During injury, some LF and Spont neurons are converted to RS. This conversion results in 77.8% RS, 22.2% LF and 0.0% Spont after injury. In the left hemisphere, PKCδ population is initialized with 70.9% RS, 19.3% LF and 9.6% Spont. During injury, some RS and Spont neurons are converted to LF and the resulted PKCδ population after injury is 46.9% RS, 46.9% LF, and 6.2% Spont.

In the right hemisphere, the CGRPR population is initialized with 58.8% RS, 35.2% LF and 5.8% Spont. During injury, some RS and Spont neurons are converted to LF. This conversion results in 50% RS, 44.4% LF and 5.6% Spont after injury. In the left hemisphere, the CGRPR population is initialized with 37.5% RS, 62.5% LF and 0% Spont. During injury, some LF neurons are converted to RS and the resulted CGRPR population after injury is 67.7% RS, 32.3% LF, and 0% Spont.

**6.5 Update neuron firing rates (Assign-Firing-Rates).** A stochastic procedure is used to update all neuron firing rates. The firing rate of a neuron is determined by stimulation intensity and the neuron's cell-type, firing rate type, and damage level.

The average firing rate for a neuron based on its cell-type, rate type, and stimulation during uninjured (control) conditions is defined as $\mu_{Control}$. Similarly, average firing rate for a neuron during injured (CYP-sensitized) conditions is defined as $\mu_{Injury}$. These values are obtained from the tables established during initialization. Separate tables are used for the left and right CeA. To determine a neuron's firing rate at time step *t*, two random variables are drawn from Poisson distributions:

- *X* is sampled from a Poisson distribution with mean $\mu_{Control}$
- *Y* is sampled from a Poisson distribution with mean $\mu_{Injury}$

At each time step, the neuron's firing rate (*FR*) is calculated as a weighted average of *X* and *Y:*

$$FR = \left(\frac{100 - d}{100}\right) \cdot X + \left(\frac{d}{100}\right) \cdot Y$$

where *d* is the neuron's damage. When *d = 0*, the neuron's firing rate is sampled from the distribution with mean $\mu_{Control}$. When *d = 100*, the neuron's firing rate is sampled from the distribution with mean $\mu_{Injury}$. When *0 < d < 100*, the neuron's firing rate is a weighted combination of these two distributions, allowing neuron firing rates to transition proportionally toward the injured state.

Spontaneous neurons are assigned constant firing rates that do not vary throughout a simulation. In the right CeA, PKCδ Spont neurons fire at 4.143 Hz and CGRPR Spont neurons at 4 Hz. In the left CeA, PKCδ Spont neurons fire at 3.25 Hz and CGRPR Spont neurons at 4 Hz.

**6.6 Send inhibitory Signals (Send-Inhibitory-Signals).** After firing rates are updated, the network is used to send inhibitory signals between connected neurons. Each neuron transmits an inhibitory signal to its connected targets based on its current firing rate. The strength of each inhibitory signal is equal to the firing rate of the transmitting neuron.

For each neuron in the network, all incoming signals are summed to calculate the total inhibitory input. If a neuron's total inhibitory input is greater than or equal to 15 Hz, the neuron is considered inhibited, and its firing rate is set to zero (*FR = 0)*. If the total input is below the threshold, the neuron's firing rate remains unchanged.

**6.7 Calculate Model Output (Calculate-Pain).** At each time step, nociceptive output from the left CeA and right CeA is calculated separately as the sum of the firing rate activity across all PKCδ and CGRPR neurons that express pERK. A measure of total nociceptive output is then calculated as the sum of nociceptive output from the left and right CeA.

In the right CeA, PKCδ and CGRPR neurons are assumed to be pro-nociceptive (increase pain). The nociceptive output from the right CeA at each time step calculated as

$$P_{Right} = \sum_{i \in PKC\delta+ \,/\, \mathrm{pERK+}} \frac{d_i}{100} \cdot FR_i + \sum_{i \in CGRPR+ \,/\, \mathrm{pERK+}} FR_i$$

The first summation in the above equation represents the contribution to nociceptive signaling from PKCδ neurons (including the co-expressing PKC$\delta$+CGRPR neurons assigned PKCδ attributes) and the second summation represents the contribution to nociceptive signaling from CGRPR neurons (including the co-expressing neurons assigned CGRPR attributes). Because the nociceptive output each PKCδ neuron is scaled by damage, these neurons start to contribute to nociceptive output when $d > 0$ and contribute fully when $d = 100$.

In the left CeA, PKC$\delta$ and CGRPR neurons are assumed to be anti-nociceptive (decrease pain). Similar to above, the nociceptive output from the left CeA at each time step is calculated as

$$P_{Left} = -\left( \sum_{i \in PKC\delta+ \,/\, \mathrm{pERK+}} \frac{d_i}{100} \cdot FR_i + \sum_{i \in CGRPR+ \,/\, \mathrm{pERK+}} FR_i \right)$$

Total nociceptive output from the bilateral CeA is then calculated as $P_{Total} = P_{Right} + P_{Left}$.

**Supplemental Figures**

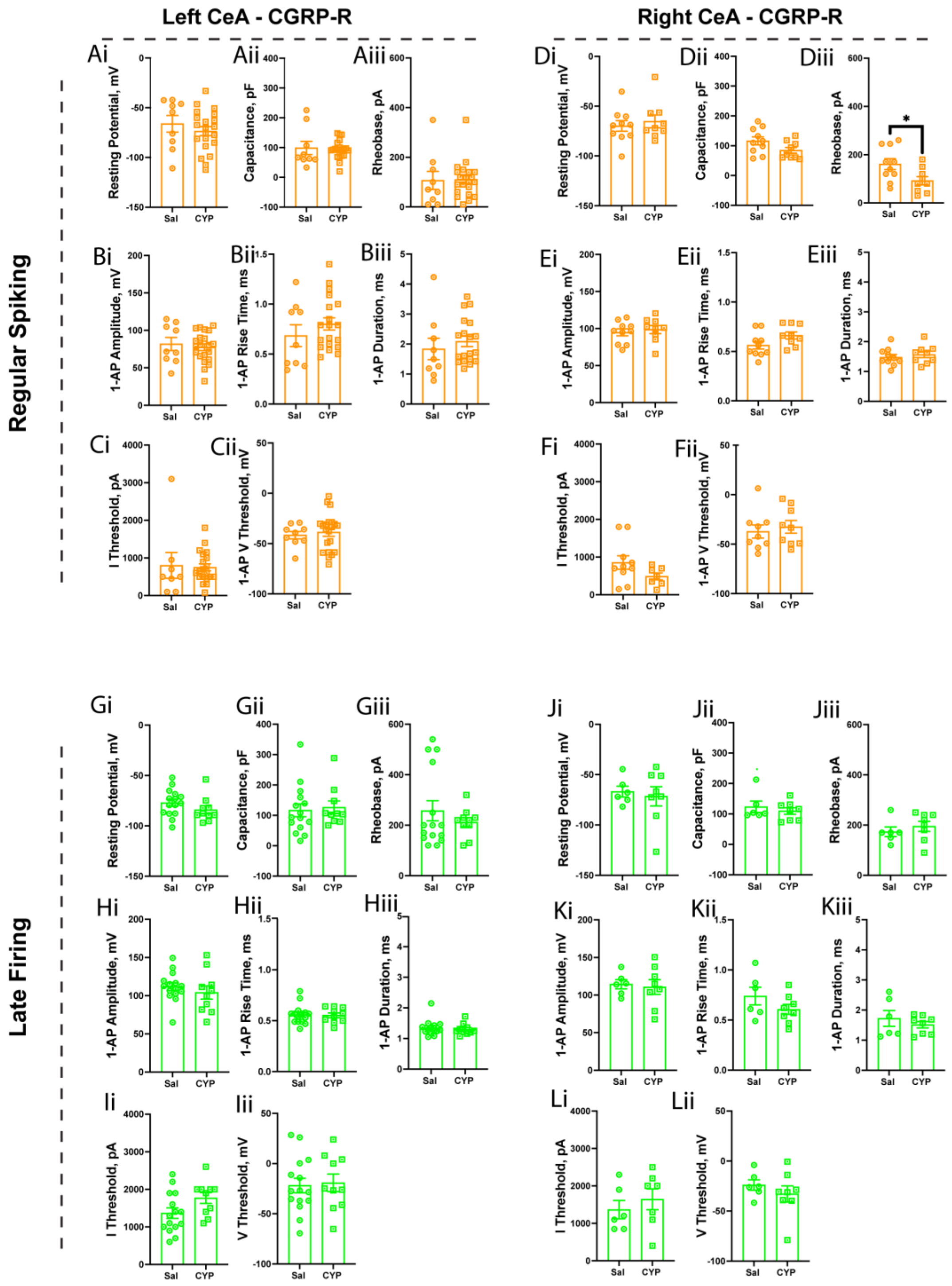


**Figure S1: Single action potential properties of CGRP-R positive neurons in the left or right CeA. (Ai-Cii)** In the regular-spiking type, no change were observed in the left CeA's single action potential properties - resting potential, capacitance, rheobase, amplitude, rise time, duration, I-Threshold and V-Threshold between saline and CYP groups. **(Di, Dii, Ei-Fii)** The right CeA's single action potential properties - resting potential, capacitance, amplitude, rise time, duration, I-Threshold and V-Threshold remained unchanged between saline and CYP groups, **(Diii)** but the rheobase was significantly reduced in the CYP group. **(Gi-Iii, Ji-Lii)** No changes were observed in the left and right CeA's late-firing neuronal single action potential properties - resting potential, capacitance, rheobase, amplitude, rise time, duration, I-Threshold and V-Threshold between saline and CYP groups. All data are presented as mean + SEM, and error bars represent SEM. Statistical test used – unpaired T-test, $*p<0.05$.

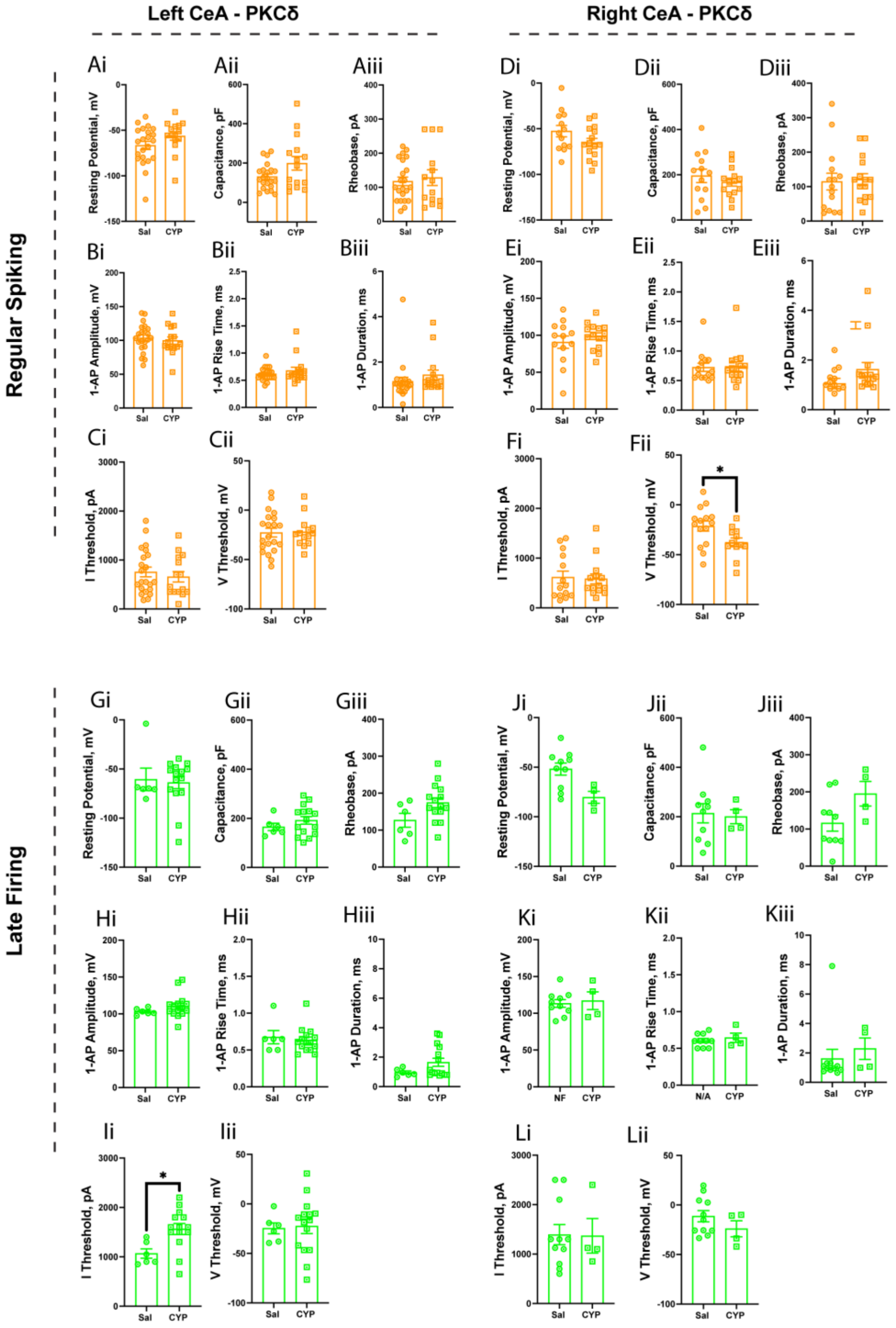


**Figure S2: Single action potential properties of PKCδ positive neurons in the left or right CeA. (Ai-Cii)** In the regular-spiking type, no changes were observed in the left CeA's single action potential properties - resting potential, capacitance, rheobase, amplitude, rise time, duration, I-Threshold and V-Threshold between saline and CYP groups. **(Di-Di, Ei-Fi)** The right CeA's single action potential properties - resting potential, capacitance, rheobase, amplitude, rise time, duration and I-Threshold remained unchanged between saline and CYP groups, **(Fii)** but the V-Threshold was significantly reduced in the CYP group. **(Gi-Hiii, Iii, Ji-Lii)** No changes were observed in the left and right CeA's late-firing neuronal single action potential properties - resting potential, capacitance, rheobase, amplitude, rise time, duration and V-Threshold between saline and CYP groups, **(Ii)** except in the current threshold where the left CeA exhibited increased current threshold indicating an reduced excitability. All data are presented as mean + SEM, and error bars represent SEM. Statistical test used – unpaired T-test, *p<0.05.

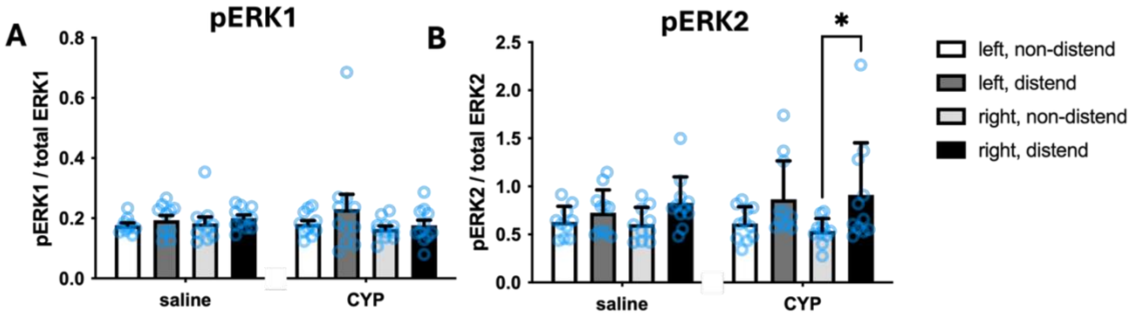


**Figure S3: Evaluation of ERK phosphorylation in the CeA.** pERK1/2 levels in the left and right CeA were evaluated via Western blot analysis and normalized to total ERK1/2 levels following UBD in saline and CYP-sensitized animals. (**A**) Graded bladder distension failed to increase pERK1 expression in saline or CYP-sensitized animals (Two-way RM ANOVA, no main effect of distension in saline P=0.2259 or CYP P=0.3161; n=10-11). (**B**) One set of graded bladder distension increased pERK2 expression in both saline and CYP-sensitized animals (Two-way RM ANOVA, main effect of distension in saline p=0.0349, main effect of distension in CYP p=0.0298; n=11). There were no side-specific increases in pERK2 expression in saline-treated animals (Bonferroni post-test distend left vs. distend right p>0.05, left sham (non-distend) vs. left distend p>0.05, right sham vs. right distend p>0.05), however bladder distension significantly increased pERK2 in the right CeA of CYP-treated mice as compared to CYP-treated sham animals (Bonferroni post-test CYP right, sham vs. distend *p<0.05). Data are mean ± SEM with individual data points plotted.

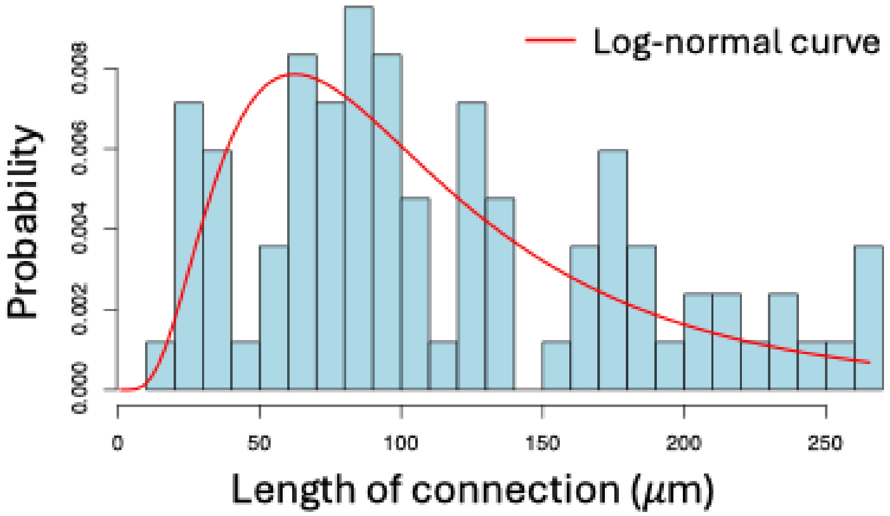


**Figure S4: Distribution of observed dendritic lengths from PKC neurons.** (**A**) Bars indicate observed lengths. The red line indicates the best fitting log-normal distribution. This fitted distribution was used to define the probabilities of short (< 30 m), medium (30 – 120 m), long (120 – 200 m), and extra-long (200 – 270 m) connections in model. N=7 total cells. Original data are from Adke et al, 2021.

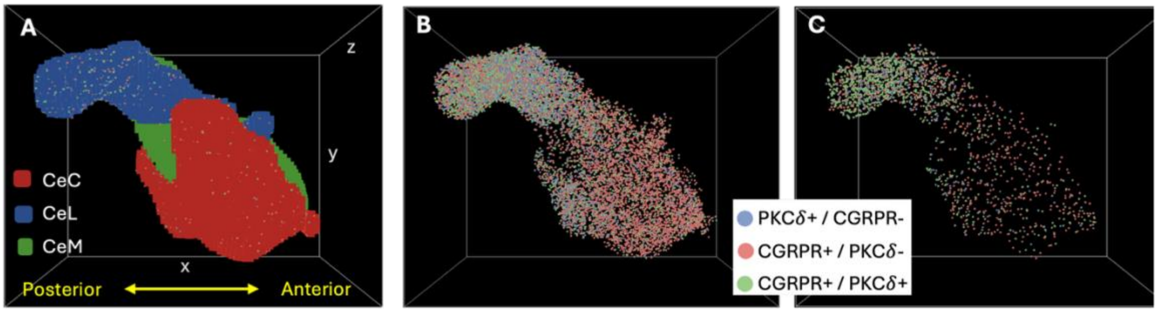


**Figure S5: NetLogo3D visualization of CeA and neuron populations: (A)** Spatial domain showing patches corresponding to the capsular (CeC = red), lateral (CeL = blue), and medial (CeM = green) subnuclei. **(B)** CeA spatial domain without patch overlay, showing an example distribution of PKC$\delta$ and CGRPR neurons. **(C)** The same distribution restricted to pERK+ neurons.

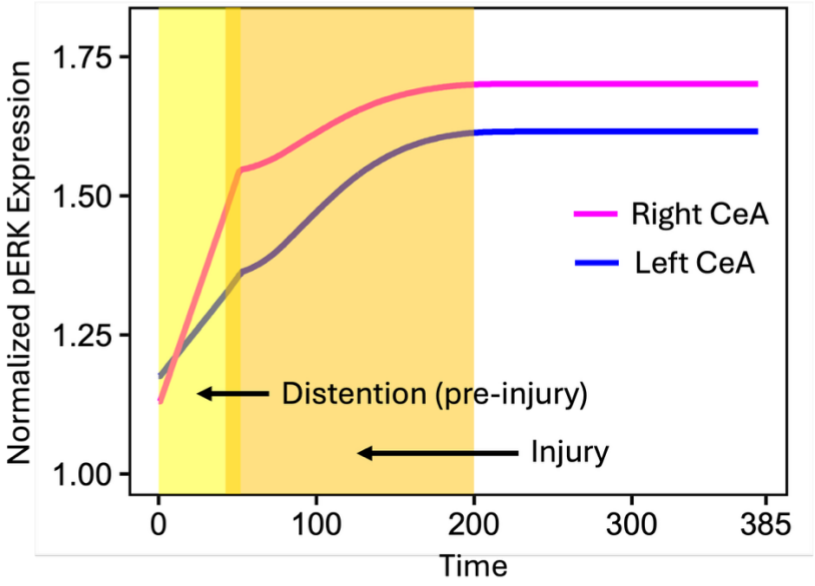


**Figure S6: Model predictions of pERK expression in the left and right CeA.** The y-axis shows the total number of pERK-positive neurons, normalized to post-injury expression levels. pERK expression increases during pre-injury distention (yellow shaded window) and further during injury (orange shaded window). Blue and pink curves denote the left and right CeA, respectively.

**Tables**

**Table S1. Agent variables assigned to each neuron in the ABM.**

| Variable | Description | Value | Frequency of updates |
|---|---|---|---|
| *xcor, ycor, zcor* | Variables indicating 3-D location of neuron within CeA | Coordinates within CeA patches | Assigned at initialization |
| *PKC?* | Expression-type status | Yes or No | Assigned at initialization |
| *CGRPR?* | Expression-type status | Yes or No | Assigned at initialization |
| *pERK?* | Expression-type status | Yes or No | Assigned at initialization and updated during simulation |
| *Rate-Type* | Firing rate pattern | Late Firing (LF), Regular Spiking (RS), or Spontaneous (Spont) | Assigned at initialization and updated during simulation |
| *Latency* | Length of damage latency period | integer in [40,80] | Assigned at initialization |
| *Sensitivity* | Length of sensitizing period | integer in [50,150] | Assigned at initialization |
| *Damage* | Damage (percent of total damage) | value in [0,100] | Updated each time step |
| *Incoming-Connections* | Number of incoming connections | Integer | Assigned at initialization |
| *Outgoing-Connections* | Number of outgoing connections | Integer | Assigned at initialization |
| *Incoming-Connections-IDs* | List of IDs of neurons sending signals to agent | List of agent IDs | Assigned at initialization |
| *Outgoing-Connections-IDs* | List of IDs of neurons that agent sends signals to | List of agent IDs | Assigned at initialization |
| *Inhibited?* | Inhibition status | Yes or No | Updated each time step |
| *Firing-Rate* | Firing rate (Hz) | non-negative value | Updated each time step |

**Table S2. Global variables in the ABM.**

| Variable | Description | Frequency of updates |
|---|---|---|
| *Stim* | Stimulation intensity | Updated each time step |
| $S_{Total}$ | Total time under stimulation | Updated each time step |
| $P_{Left}$ | Pain output from left CeA | Updated each time step |
| $P_{Right}$ | Pain output from right CeA | Updated each time step |
| $P_{Total}$ | Total pain output | Updated each time step |
| *Avg-Damage* | Average damage across all neurons | Updated each time step |

**Table S3. Parameters defining the Poisson distributions for neuron firing rates.** Firing rates are drawn from a Poisson distribution with mean $\mu$ obtained from experimental data. Mean values vary based on hemisphere (left or right), control or CYP-sensitized (injured) condition, cell-type (PKC$\delta$ or CGRPR) and firing frequency (late firing - LF or regular spiking - RS), and stimulation intensity (0 to 360 pA).

| Cell Type | Hemisphere | Firing Freq. | Stimulation (pA) | Control | CYP-sensitized |
|---|---|---|---|---|---|
| | | | | $\mu$ | $\mu$ |
| PKC$\delta$ | Right | LF | 0-40 | 0 | 0 |
| PKC$\delta$ | Right | LF | 80 | 0.125 | 0 |
| PKC$\delta$ | Right | LF | 120 | 1.375 | 0 |
| PKC$\delta$ | Right | LF | 160 | 3.875 | 0.33333333 |
| PKC$\delta$ | Right | LF | 200 | 6.125 | 2 |
| PKC$\delta$ | Right | LF | 240 | 7.5 | 3.66666667 |
| PKC$\delta$ | Right | LF | 280 | 8.125 | 5.66666667 |
| PKC$\delta$ | Right | LF | 320 | 8.875 | 6.66666667 |
| PKC$\delta$ | Right | LF | 360 | 9.625 | 6.33333333 |
| PKC$\delta$ | Right | RS | 0 | 0 | 0 |
| PKC$\delta$ | Right | RS | 40 | 0.8 | 0 |
| PKC$\delta$ | Right | RS | 80 | 2.2 | 0.71428571 |
| PKC$\delta$ | Right | RS | 120 | 2.9 | 2.71428571 |
| PKC$\delta$ | Right | RS | 160 | 4 | 6 |
| PKC$\delta$ | Right | RS | 200 | 5.15 | 8.42857143 |
| PKC$\delta$ | Right | RS | 240 | 6.45 | 10.3571429 |
| PKC$\delta$ | Right | RS | 280 | 7.45 | 10.7857143 |
| PKC$\delta$ | Right | RS | 320 | 7.75 | 10.2142857 |
| PKC$\delta$ | Right | RS | 360 | 8.35 | 10.3571429 |
| PKC$\delta$ | Left | LF | 0-40 | 0 | 0 |
| PKC$\delta$ | Left | LF | 80 | 1.2 | 0.07692308 |
| PKC$\delta$ | Left | LF | 120 | 4 | 0.61538462 |
| PKC$\delta$ | Left | LF | 160 | 6.4 | 1.53846154 |
| PKC$\delta$ | Left | LF | 200 | 9.2 | 3.07692308 |
| PKC$\delta$ | Left | LF | 240 | 11 | 5 |
| PKC$\delta$ | Left | LF | 280 | 12.6 | 6.23076923 |
| PKC$\delta$ | Left | LF | 320 | 13.2 | 7.23076923 |
| PKC$\delta$ | Left | LF | 360 | 12.8 | 7.76923077 |
| PKC$\delta$ | Left | RS | 0-40 | 0 | 0.0625 |
| PKC$\delta$ | Left | RS | 80 | 1.30434783 | 1.8125 |
| PKC$\delta$ | Left | RS | 120 | 3.82608696 | 3.3125 |
| PKC$\delta$ | Left | RS | 160 | 5.91304348 | 5.5 |
| PKC$\delta$ | Left | RS | 200 | 7.73913043 | 7.3125 |
| PKC$\delta$ | Left | RS | 240 | 9.60869565 | 8.4375 |
| PKC$\delta$ | Left | RS | 280 | 11.0434783 | 9 |
| PKC$\delta$ | Left | RS | 320 | 11.3043478 | 9.3125 |
| PKC$\delta$ | Left | RS | 360 | 11.3043478 | 9.0625 |
| CGRPR | Right | LF | 0 | 0 | 0 |
| CGRPR | Right | LF | 40 | 0 | 0.25 |
| CGRPR | Right | LF | 80 | 0 | 1.25 |
| CGRPR | Right | LF | 120 | 0 | 2 |
| CGRPR | Right | LF | 160 | 0 | 2.75 |
| CGRPR | Right | LF | 200 | 2 | 3.75 |
| CGRPR | Right | LF | 240 | 3.5 | 5.5 |

| | | | | | |
|---|---|---|---|---|---|
| CGRPR | Right | LF | 280 | 4 | 7.5 |
| CGRPR | Right | LF | 320 | 5 | 8 |
| CGRPR | Right | LF | 360 | 5.5 | 8.5 |
| CGRPR | Right | RS | 0 | 0 | 0 |
| CGRPR | Right | RS | 40 | 0 | 0.2 |
| CGRPR | Right | RS | 80 | 0.07142857 | 1.2 |
| CGRPR | Right | RS | 120 | 0.71428571 | 2.7 |
| CGRPR | Right | RS | 160 | 2 | 4.4 |
| CGRPR | Right | RS | 200 | 3.28571429 | 5.9 |
| CGRPR | Right | RS | 240 | 4.28571429 | 7.5 |
| CGRPR | Right | RS | 280 | 5.28571429 | 8.5 |
| CGRPR | Right | RS | 320 | 6.14285714 | 8.9 |
| CGRPR | Right | RS | 360 | 6.35714286 | 9.6 |
| CGRPR | Left | LF | 0 | 0 | 0 |
| CGRPR | Left | LF | 40 | 0 | 0 |
| CGRPR | Left | LF | 80 | 0 | 0 |
| CGRPR | Left | LF | 120 | 0 | 0.8 |
| CGRPR | Left | LF | 160 | 0 | 2 |
| CGRPR | Left | LF | 200 | 0.33333333 | 2.6 |
| CGRPR | Left | LF | 240 | 1 | 4 |
| CGRPR | Left | LF | 280 | 2 | 6.4 |
| CGRPR | Left | LF | 320 | 3.33333333 | 8 |
| CGRPR | Left | LF | 360 | 4 | 9.8 |
| CGRPR | Left | RS | 0 | 0 | 0 |
| CGRPR | Left | RS | 40 | 1.19047619 | 0.31818182 |
| CGRPR | Left | RS | 80 | 2.14285714 | 0.5 |
| CGRPR | Left | RS | 120 | 2.33333333 | 1.22727273 |
| CGRPR | Left | RS | 160 | 3.04761905 | 2.19047619 |
| CGRPR | Left | RS | 200 | 4.42857143 | 4.22727273 |
| CGRPR | Left | RS | 240 | 5.66666667 | 5.31818182 |
| CGRPR | Left | RS | 280 | 6.76190476 | 6.09090909 |
| CGRPR | Left | RS | 320 | 7.42857143 | 6.40909091 |
| CGRPR | Left | RS | 360 | 8 | 6.86363636 |

**Table S4. Statistical Details for Electrophysiology Experiments**

| Figure | Normality | Test | Sample Size | Data |
|---|---|---|---|---|
| Figure 1E | No assumptions made | Fisher's Exact | Sal - LF-15, RS-9, spon-0<br>CYP - LF-10, 21, spon-0 | p=0.0321 (two-sided) |
| Figure 1F | No assumptions made | Fisher's Exact | Sal - LF-6, RS-10, spon-1<br>CYP - LF-8, 9, spon-0 | p=0.8603 (two-sided) |
| Figure 1H | No assumptions made | Fisher's Exact | Sal - Acc-14, NA-10<br>CYP - Acc-13, NA-21 | p=0.1828 (two-sided) |
| Figure 1I | No assumptions made | Fisher's Exact | Sal - Acc-6, NA-11<br>CYP - Acc-8, NA-10 | p=0.7332 (two-sided) |
| Figure 1 K1 | Two independent variables:<br>1. Injury State,<br>2. Current Injection | Two-way ANOVA | Sal - 9<br>CYP - 21 | Injury Status: F (1, 279) = 14.83; p=0.0001<br>Current Injection: F (9, 279) = 7.266; p<0.0001 |
| Figure 1 K2 | Non-normal distribution | Mann-Whitney *U* test | Sal - 9<br>CYP - 21 | p=0.0718 (two-tailed); *U*=56 |
| Figure 1 M1 | Two independent variables: 1. Injury State, 2. Current Injection | Two-way ANOVA | Sal - 10<br>CYP - 9 | Injury Status: F (1, 170) = 33.55; p<0.0001<br>Current Injection: F (9, 170) = 18.42; p<0.0001 |
| Figure 1 M2 | Non-normal distribution | Mann-Whitney *U* test | Sal - 10<br>CYP - 9 | p=0.0163 (two-tailed); *U*=17.50 |
| Figure 1 O1 | Two independent variables: 1. Injury State, 2. Current Injection | Two-way ANOVA | Sal - 15<br>CYP - 10 | Injury Status: F (1, 230) = 0.5564; p=0.4565<br>Current Injection: F (9, 230) = 25.01; p<0.0001 |
| Figure 1 O2 | Non-normal distribution | Mann-Whitney *U* test | Sal - 15<br>CYP -10 | p=0.6673 (two-tailed); *U*=67 |
| Figure 1 Q1 | Two independent variables: 1. Injury State, 2. Current Injection | Two-way ANOVA | Sal - 6<br>CYP - 8 | Injury Status: F (1, 120) = 16.17; p=0.0001<br>Current Injection: F (9, 120) = 19.25; p<0.0001 |
| Figure 1 Q2 | Non-normal distribution | Mann-Whitney *U* test | Sal - 6<br>CYP -8 | p=0.0053 (two-tailed); *U*=3 |
| Figure 2E | No assumptions made | Fisher's Exact | Sal - LF-6, RS-22, spon-3<br>CYP - LF-15, 15, spon-2 | p=0.0766 (two-sided) |
| Figure 2F | No assumptions made | Fisher's Exact | Sal - LF-11, RS-17, spon-7<br>CYP - LF-4, RS-14, spon-0 | p=0.0543 (two-sided) |
| Figure 2H | No assumptions made | Fisher's Exact test | Sal - Acc-12, NA-16<br>CYP - Acc-15, NA-15 | p=0.6096 (two-sided) |

| | | | | |
|---|---|---|---|---|
| Figure 2I | No assumptions made | Fisher's Exact test | Sal - Acc-12, NA-16<br>CYP - Acc-6, NA-12 | p=0.5539 (two-sided) |
| Figure 2 K1 | Two independent variables: 1. Injury State, 2. Current Injection | Two-way ANOVA | Sal - 22<br>CYP - 15 | Injury Status: F (1, 350) = 3.088; p=0.0797<br>Current Injection: F (9, 350) = 22.18; p<0.0001 |
| Figure 2 K2 | Non-normal distribution | Mann-Whitney *U* test | Sal - 22<br>CYP - 15 | p=0.6733 (two-tailed); *U*=151 |
| Figure 2 M1 | Two independent variables: 1. Injury State, 2. Current Injection | Two-way ANOVA | Sal - 17<br>CYP - 14 | Injury Status: F (1, 290) = 6.191; p=0.0134<br>Current Injection: F (9, 290) = 19.52; p<0.0001 |
| Figure 2 M2 | Non-normal distribution | Mann-Whitney *U* test | Sal - 17<br>CYP - 14 | p=0.1290 (two-tailed); *U*=80.50 |
| Figure 2 O1 | Two independent variables: 1. Injury State, 2. Current Injection | Two-way ANOVA | Sal - 6<br>CYP - 15 | Injury Status: F (1, 190) = 45.44; p<0.0001<br>Current Injection: F (9, 190) = 28.88; p<0.0001 |
| Figure 2 O2 | Normal distribution, Unequal variance | Welch's *t* test | Sal - 6<br>CYP - 15 | p=0.1662; t=1.593, df=5.544 (two-tailed) |
| Figure 2 Q1 | Two independent variables: 1. Injury State, 2. Current Injection | Two-way ANOVA | Sal - 11<br>CYP - 4 | Injury Status: F (1, 130) = 10.82; p=0.0001<br>Current Injection: F (9, 130) = 9.173; p<0.0001 |
| Figure 2 Q2 | Normal distribution, Same variance | Unpaired *t* test | Sal - 11<br>CYP - 4 | p=0.1737; t=1.439, df=13 (two-tailed) |
| Figure S1 Ai | Normal distribution, Same variance | Unpaired *t* test, | Sal - 9<br>CYP - 21 | p=0.4127; t=0.8315, df=28 (two-tailed) |
| Figure S1 Aii | Non-normal distribution | Mann-Whitney *U* test | Sal - 9<br>CYP - 21 | p=0.3722 (two-tailed); *U*=74 |
| Figure S1 Aiii | Non-normal distribution | Mann-Whitney *U* test | Sal - 9<br>CYP - 21 | p=0.5852 (two-tailed); *U*=82 |
| Figure S1 Bi | Non-normal distribution | Mann-Whitney *U* test | Sal - 9<br>CYP - 21 | p=0.6892 (two-tailed); *U*=85 |
| Figure S1 Bii | Normal distribution, Same variance | Unpaired *t* test, | Sal - 9<br>CYP - 21 | p=0.3222; t=1.009, df=26 (two-tailed) |
| Figure S1 Biii | Non-normal distribution | Mann-Whitney *U* test | Sal - 9<br>CYP - 21 | p=0.2281 (two-tailed); *U*=60.50 |
| Figure S1 Ci | Non-normal distribution | Mann-Whitney *U* test | Sal - 9<br>CYP - 21 | p=0.4004 (two-tailed); *U*=63 |
| Figure S1 Cii | Normal distribution Same variance | Unpaired *t* test, | Sal - 9<br>CYP - 21 | p=0.6676; t=0.4340, df=28 (two-tailed) |

| | | | | |
|---|---|---|---|---|
| Figure S1 Di | Non-normal distribution | Mann-Whitney *U* test | Sal - 10<br>CYP - 9 | p=0.7802 (two-tailed); *U*=41 |
| Figure S1 Dii | Normal distribution, Same variance | Unpaired *t* test, | Sal - 10<br>CYP - 9 | p=0.0783; t=1.874, df=17 (two-tailed) |
| Figure S1 Diii | Normal distribution, Same variance | Unpaired *t* test, | Sal - 10<br>CYP - 9 | p=0.0272; t=2.417, df=17 (two-tailed) |
| Figure S1 Ei | Normal distribution, Same variance | Unpaired *t* test, | Sal - 10<br>CYP - 9 | p=0.5976; t=0.5379, df=17 (two-tailed) |
| Figure S1 Eii | Normal distribution, Same variance | Unpaired *t* test, | Sal - 10<br>CYP - 9 | p=0.0786; t=1.871, df=17 (two-tailed) |
| Figure S1 Eiii | Normal distribution, Same variance | Unpaired *t* test, | Sal - 10<br>CYP - 9 | p=0.5166; t=0.6624, df=17 (two-tailed) |
| Figure S1 Fi | Normal distribution, Same variance | Unpaired *t* test, | Sal - 10<br>CYP - 9 | p=0.1287; t=1.608, df=15 (two-tailed) |
| Figure S1 Fii | Normal distribution, Same variance | Unpaired *t* test, | Sal - 10<br>CYP - 9 | p=0.5061; t=0.6802, df=16 (two-tailed) |
| Figure S1 Gi | Non-normal distribution | Mann-Whitney *U* test | Sal - 15<br>CYP - 10 | p=0.0910 (two-tailed); *U*=44 |
| Figure S1 Gii | Non-normal distribution | Mann-Whitney *U* test | Sal - 15<br>CYP - 10 | p=0.5671 (two-tailed); *U*=64 |
| Figure S1 Giii | Non-normal distribution | Mann-Whitney *U* test | Sal - 15<br>CYP - 10 | p=0.7324 (two-tailed); *U*=68.50 |
| Figure S1 Hi | Normal distribution, Same variance | Unpaired *t* test, | Sal - 15<br>CYP - 10 | p=0.4153; t=0.8295, df=23 (two-tailed) |
| Figure S1 Hii | Normal distribution, Same variance | Unpaired *t* test, | Sal - 15<br>CYP - 10 | p=0.8448; t=0.1979, df=23 (two-tailed) |
| Figure S1 Hiii | Non-normal distribution | Mann-Whitney *U* test | Sal - 15<br>CYP - 10 | p=0.6357 (two-tailed); *U*=61.50 |
| Figure S1 Ii | Normal distribution, Same variance | Unpaired *t* test, | Sal - 15<br>CYP - 10 | p=0.0639; t=1.946, df=23 (two-tailed) |
| Figure S1 Iii | Normal distribution, Same variance | Unpaired *t* test, | Sal - 15<br>CYP - 10 | p=0.8172; t=0.2338, df=23 (two-tailed) |
| Figure S1 Ji | Non-normal distribution | Mann-Whitney *U* test | Sal - 6<br>CYP - 8 | p=0.9497 (two-tailed); *U*=23 |
| Figure S1 Jii | Non-normal distribution | Mann-Whitney *U* test | Sal - 6<br>CYP - 8 | p=0.7546 (two-tailed); *U*=21 |
| Figure S1 Jiii | Non-normal distribution | Mann-Whitney *U* test | Sal - 6<br>CYP - 8 | p=0.3919 (two-tailed); *U*=17 |
| Figure S1 Ki | Normal distribution, Same variance | Unpaired *t* test, | Sal - 6<br>CYP - 8 | p=0.7728; t=0.2953, df=12 (two-tailed) |
| Figure S1 Kii | Normal distribution, Same variance | Unpaired *t* test, | Sal - 6<br>CYP - 8 | p=0.1881; t=1.396, df=12 (two-tailed) |

| | | | | |
|---|---|---|---|---|
| Figure S1 Kiii | Non-normal distribution | Mann-Whitney *U* test | Sal - 6<br>CYP - 8 | p=0.5961 (two-tailed); *U*=19.50 |
| Figure S1 Li | Normal distribution, Same variance | Unpaired *t* test, | Sal - 6<br>CYP - 8 | p=0.4789; t=0.7330, df=11 (two-tailed) |
| Figure S1 Lii | Normal distribution, Same variance | Unpaired *t* test, | Sal - 6<br>CYP - 8 | p=0.8738; t=0.1622, df=12 (two-tailed) |
| Figure S2 Ai | Non-normal distribution | Mann-Whitney *U* test | Sal - 22<br>CYP - 15 | p=0.1049 (two-tailed); *U*=112 |
| Figure S2 Aii | Normal distribution, Same variance | Unpaired *t* test, | Sal - 22<br>CYP - 15 | p=0.0457; t=2.072, df=35 (two-tailed) |
| Figure S2 Aiii | Non-normal distribution | Mann-Whitney *U* test | Sal - 22<br>CYP - 15 | p=0.9808 (two-tailed); *U*=153 |
| Figure S2 Bi | Normal distribution, Same variance | Unpaired *t* test, | Sal - 22<br>CYP - 15 | p=0.5737; t=0.5680, df=35 (two-tailed) |
| Figure S2 Bii | Non-normal distribution | Mann-Whitney *U* test | Sal - 22<br>CYP - 15 | p=0.6977 (two-tailed); *U*=131 |
| Figure S2 Biii | Non-normal distribution | Mann-Whitney *U* test | Sal - 22<br>CYP - 15 | p=0.1322 (two-tailed); *U*=116 |
| Figure S2 Ci | Non-normal distribution | Mann-Whitney *U* test | Sal - 22<br>CYP - 15 | p=0.4855 (two-tailed); *U*=142 |
| Figure S2 Cii | Non-normal distribution | Mann-Whitney *U* test | Sal - 22<br>CYP - 15 | p=0.8812 (two-tailed); *U*=142 |
| Figure S2 Di | Normal distribution, Same variance | Unpaired *t* test, | Sal - 13<br>CYP - 15 | p=0.1094; t=1.658, df=26 (two-tailed) |
| Figure S2 Dii | Non-normal distribution | Mann-Whitney *U* test | Sal - 13<br>CYP - 15 | p=0.4879 (two-tailed); *U*=76 |
| Figure S2 Diii | Non-normal distribution | Mann-Whitney *U* test | Sal - 13<br>CYP - 15 | p=0.5722 (two-tailed); *U*=98.50 |
| Figure S2 Ei | Normal distribution, Same variance | Unpaired *t* test, | Sal - 13<br>CYP - 15 | p=0.3940; t=0.8667, df=26 (two-tailed) |
| Figure S2 Eii | Non-normal distribution | Mann-Whitney *U* test | Sal - 13<br>CYP - 15 | p=0.8042 (two-tailed); *U*=99 |
| Figure S2 Eiii | Non-normal distribution | Mann-Whitney *U* test | Sal - 13<br>CYP - 15 | p=0.0819 (two-tailed); *U*=59.50 |
| Figure S2 Fi | Non-normal distribution | Mann-Whitney *U* test | Sal - 13<br>CYP - 15 | p=0.7082 (two-tailed); *U*=89.50 |
| Figure S2 Fii | Normal distribution, Same variance | Unpaired *t* test, | Sal - 13<br>CYP - 15 | p=0.0256; t=2.374, df=25 (two-tailed) |
| Figure S2 Gi | Normal distribution, Same variance | Unpaired *t* test, | Sal - 6<br>CYP - 15 | p=0.02947; t=1.078, df=19 (two-tailed) |

| | | | | |
|---|---|---|---|---|
| Figure S2 Gii | Normal distribution, Same variance | Unpaired *t* test, | Sal - 6<br>CYP - 15 | p=0.3337; t=0.9920, df=19 (two-tailed) |
| Figure S2 Giii | Normal distribution, Same variance | Unpaired *t* test, | Sal - 6<br>CYP - 15 | p=0.0603; t=1.997, df=19 (two-tailed) |
| Figure S2 Hi | Normal distribution, Same variance | Unpaired *t* test, | Sal - 6<br>CYP - 15 | p=0.2947; t=1.078, df=19 (two-tailed) |
| Figure S2 Hii | Non-normal distribution | Mann-Whitney *U* test | Sal - 6<br>CYP - 15 | p=0.8024 (two-tailed); *U*=41.50 |
| Figure S2 Hiii | Non-normal distribution | Mann-Whitney *U* test | Sal - 6<br>CYP - 15 | p=0.2910 (two-tailed); *U*=31 |
| Figure S2 Ii | Normal distribution, Same variance | Unpaired *t* test, | Sal - 6<br>CYP - 15 | p=0.0135; t=2.739, df=18 (two-tailed) |
| Figure S2 Iii | Normal distribution, Same variance | Unpaired *t* test, | Sal - 6<br>CYP - 15 | p=0.2526; t=1.180, df=19 (two-tailed) |
| Figure S2 Ji | Normal distribution, Same variance | Unpaired *t* test, | Sal - 10<br>CYP - 4 | p=0.0.0192; t=2.704, df=12 (two-tailed) |
| Figure S2 Jii | Normal distribution, Same variance | Unpaired *t* test, | Sal - 10<br>CYP - 4 | p=0.8360; t=0.2116, df=12 (two-tailed) |
| Figure S2 Jiii | Normal distribution, Same variance | Unpaired *t* test, | Sal - 10<br>CYP - 4 | p=0.0765; t=1.938, df=12 (two-tailed) |
| Figure S2 Ki | Normal distribution, Same variance | Unpaired *t* test, | Sal - 10<br>CYP - 4 | p=0.7477; t=0.3291, df=12 (two-tailed) |
| Figure S2 Kii | Normal distribution, Same variance | Unpaired *t* test, | Sal - 10<br>CYP - 4 | p=0.5110; t=0.6774, df=12 (two-tailed) |
| Figure S2 Kiii | Non-normal distribution | Mann-Whitney *U* test | Sal - 10<br>CYP - 4 | p=0.2125 (two-tailed); *U*=12 |
| Figure S2 Li | Normal distribution, Same variance | Unpaired *t* test, | Sal - 10<br>CYP - 4 | p=0.9521; t=0.06129, df=13 (two-tailed) |
| Figure S2 Lii | Normal distribution, Same variance | Unpaired *t* test, | Sal - 10<br>CYP - 4 | p=0.3121; t=1.055, df=12 (two-tailed) |

**Table S5: Details Effect Size Calculations from Published Wet Lab Experiments and ABM Simulations.** Shown are the calculated Hedges' *g* values for comparisons between simulated experiments and published wet-lab experiments from Allen et al, 2023. In simulation experiments, n=8 indicates that each representative mouse is simulated once, while n=48 indicates that each representative mouse was simulated 6 times. A pressure of 30mmHg represents a low noxious bladder distension stimulation in mice and is replicated by a current of 120pA in model simulations. A pressure of 60mmHg is a highly noxious bladder distention stimulation in mice and is represented as a current of 240pA in model simulations. Units for wet-lab data is Vsec. Units for simulations are "nociceptive output units" based on ABM calculation (see above).

| | | | Values for first group in comparison | | | Values for second group in comparison | | | Hedges' *g* calculated for comparison | | |
|---|---|---|---|---|---|---|---|---|---|---|---|
| **Comparison** | **Simulation or Wet-Lab** | **Stimulation Intensity** | **mean** | **SD** | **N** | **mean** | **SD** | **N** | **Hedges' g** | **CI Lower** | **CI Upper** |
| Injured Intact vs Naive Intact | Simulation | 120pA | -1837.15 | 1071.34 | 48 | 3060.21 | 1907.14 | 48 | 3.1409 | 2.5451 | 3.7366 |
| Injured Intact vs Naive Intact | Wet-Lab | 30mmHg | 2.52 | 1.48 | 30 | 2.98 | 1.54 | 65 | 0.2974 | -0.1338 | 0.7286 |
| Injured Intact vs Left CGRP Activate | Simulation | 120pA | 3086.63 | 1896.22 | 8 | -27695.125 | 16784.34 | 8 | -2.4366 | -3.6901 | -1.1832 |
| Injured Intact vs Left CGRP Activate | Wet-Lab - Optogenetic | 30mmHg | 4.00 | 3.57 | 10 | 1.67 | 0.98 | 11 | -0.8766 | -1.7404 | -0.0128 |
| Injured Intact vs Left CGRP Activate | Wet-Lab - Pharmacological | 30mmHg | 3.25 | 1.27 | 10 | 0.87 | 0.60 | 10 | -2.2879 | -3.3868 | -1.1891 |
| Injured Intact vs Right CGRP Activate | Simulation | 120pA | 3086.63 | 1896.22 | 8 | 31569 | 19381.16 | 8 | 1.9556 | 0.8078 | 3.1035 |
| Injured Intact vs Right CGRP Activate | Wet-Lab - Optogenetic | 30mmHg | 4.40 | 2.18 | 8 | 5.63 | 2.88 | 10 | 0.4506 | -0.447 | 1.3481 |
| Injured Intact vs Right CGRP Activate | Wet-Lab - Pharmacological | 30mmHg | 3.87 | 1.64 | 8 | 6.57 | 2.76 | 7 | 1.137 | 0.0992 | 2.1748 |
| Injured Intact vs Naive Intact | Simulation | 240pA | -1098.46 | 694.14 | 48 | 5299.23 | 3297.65 | 48 | 2.6634 | 2.1162 | 3.2106 |
| Injured Intact vs Naive Intact | Wet-Lab | 60mmHg | 3.78 | 1.34 | 30 | 4.73 | 1.77 | 67 | 0.5674 | 0.1329 | 1.002 |
| Injured Intact vs Left CGRP Activate | Simulation | 240pA | 5356.88 | 3335.09 | 8 | -16915.70 | 9918.46 | 8 | -2.8459 | -4.199 | -1.4929 |
| Injured intact vs Left CGRP Activate | Wet-Lab - Optogenetic | 60mmHg | 5.56 | 2.99 | 10 | 2.58 | 2.58 | 10 | -1.0225 | -1.9198 | -0.1252 |
| Injured Intact vs Left CGRP Activate | Wet-Lab - Pharmacological | 60mmHg | 3.85 | 1.14 | 10 | 1.32 | 0.51 | 10 | -2.7302 | -3.9221 | -1.5383 |
| Injured Intact vs Right CGRP Activate | Simulation | 240pA | 5356.87 | 3335.09 | 8 | 23590.50 | 14291.59 | 8 | 1.6612 | 0.5705 | 2.752 |
| Injured Intact vs Right CGRP Activate | Wet-Lab - Optogenetic | 60mmHg | 4.50 | 1.68 | 8 | 7.44 | 2.45 | 10 | 1.3034 | 0.3209 | 2.2859 |
| Injured Intact vs Right CGRP Activate | Wet-Lab - Pharmacological | 60mmHg | 4.99 | 1.30 | 8 | 8.02 | 2.07 | 8 | 1.6574 | 0.5674 | 2.7475 |
| Injured intact vs Left CGRP Inhibit | Simulation | 120pA | 3086.63 | 1896.22 | 8 | 6087 | 3777.46 | 8 | 0.9491 | -0.034 | 1.9323 |
| Injured intact vs Left CGRP Inhibit | Wet-Lab - Optogenetic | 30mmHg | 4.00 | 3.57 | 10 | 6.70 | 2.89 | 7 | 0.7721 | -0.1807 | 1.7249 |
| Injured intact vs Left CGRP Inhibit | Wet-Lab - Pharmacological | 30mmHg | 3.25 | 1.27 | 10 | 6.98 | 2.58 | 9 | 1.7878 | 0.7568 | 2.8189 |

| | | | | | | | | | | | |
|---|---|---|---|---|---|---|---|---|---|---|---|
| Injured intact vs Right CGRP Inhibit | Simulation | 120pA | 3086.63 | 1896.23 | 8 | -2838.63 | 1704.66 | 8 | -3.1071 | -4.5274 | -1.6867 |
| Injured intact vs Right CGRP Inhibit | Wet-Lab - Optogenetic | 30mmHg | 4.40 | 2.18 | 8 | 0.81 | 0.69 | 7 | -2.0241 | -3.2224 | -0.8257 |
| Injured intact vs Right CGRP Inhibit | Wet-Lab - Pharmacological | 30mmHg | 3.87 | 1.64 | 8 | 0.62 | 0.84 | 6 | -2.2245 | -3.5132 | -0.9357 |
| Injured intact vs Left CGRP Inhibit | Simulation | 240pA | 5356.88 | 3335.09 | 8 | 15875.50 | 9695.61 | 8 | 1.3717 | 0.3304 | 2.413 |
| Injured intact vs Left CGRP Inhibit | Wet-Lab - Optogenetic | 60mmHg | 5.56 | 2.99 | 10 | 6.98 | 2.58 | 7 | 0.5837 | -0.3538 | 1.5212 |
| Injured intact vs Left CGRP Inhibit | Wet-Lab - Pharmacological | 60mmHg | 3.85 | 1.14 | 10 | 7.42 | 1.73 | 9 | 2.3587 | 1.2175 | 3.5 |
| Injured intact vs Right CGRP Inhibit | Simulation | 240pA | 5356.88 | 3335.09 | 8 | -9421.13 | 5564.03 | 8 | -3.046 | -4.4503 | -1.6416 |
| Injured intact vs Right CGRP Inhibit | Wet-Lab - Optogenetic | 60mmHg | 4.50 | 1.69 | 8 | 2.10 | 2.40 | 7 | -1.1048 | -2.1381 | -0.0715 |
| Injured intact vs Right CGRP Inhibit | Wet-Lab - Pharmacological | 60mmHg | 4.99 | 1.30 | 8 | 0.43 | 0.30 | 6 | -4.2339 | -6.0889 | -2.3788 |
